# Systematic comparison of gliflozins identifies a selective anti-calcifying effect of canagliflozin beyond canonical SGLT2 inhibition

**DOI:** 10.64898/2026.11.05.764503

**Authors:** Fernandez Lucie, Brunet Manon, Ballongue Emma, Ducos Victor, Buléon Marie, Feuillet Guylène, Galaricha Eduardo, Mairal Aline, Arnaud Del Bello, Augé Nathalie, Gayrard Nathalie, Laget Jonas, Fabrice Raynaud, Diaz-Tocados Juan Miguel, Thomas Arthur, Chabbert Mickael, Arefin Samsul, Kublickiene Karolina, Stenvinkel Peter, Joost P. Schanstra, Klein Julie

**Author notes:** **Correspondence to**; UMR1297 Inserm, Institute of metabolic and cardiovascular disease (I2MC), 1 Avenue Jean Poulhes, 31432 Toulouse, France.

## Abstract

**Background:** Sodium-glucose cotransporter 2 inhibitors (SGLT2i) provide important cardiorenal benefits in chronic kidney disease (CKD), but their effects on vascular calcification remain conflicting. Whether potential anti-calcifying effects are shared across gliflozins or involve mechanisms beyond canonical SGLT2 inhibition is unknown.

**Methods:** We systematically compared canagliflozin, dapagliflozin and empagliflozin in a high-phosphate diet-induced mouse model of vascular calcification and complementary *ex vivo* and *in vitro* models, including rodent and human vascular tissues and vascular smooth muscle cells (VSMCs). Aortic transcriptional response was investigated by RNA sequencing. VSMC calcification, proliferation, migration, viability, glucose uptake and bioenergetics were investigated using complementary cellular assays, including OsteoSense staining, Annexin V/propidium iodide staining, radiolabeled 2-deoxyglucose uptake and Seahorse ATP rate analysis. SGLT2/1 expression was examined by RT-qPCR, Western blotting and immunofluorescence. We also evaluated sotagliflozin, a dual SGLT1/2 inhibitor, and mizagliflozin, a selective SGLT1 inhibitor.

**Results:** Among the three gliflozins, only canagliflozin significantly reduced aortic calcification *in vivo*. Its anti-calcifying effect was also observed in rodent aortic rings, human vascular explants, cultured VSMCs and aortic rings from diabetic db/db mice. Aortic RNA-seq revealed that canagliflozin partially reversed HPD-induced transcriptional alterations, including changes in pro-calcific signaling and cellular metabolism, while some endothelial and inflammatory responses were shared among gliflozins. In mouse VSMCs, canagliflozin reduced calcification across all tested concentrations, dose-dependently decreased osteogenic marker expression, and reduced proliferation and migration. Increased apoptotic and damaged cell fractions were observed at higher concentrations, but not at lower concentrations. Canagliflozin also reduced glucose uptake and mitochondrial ATP production, without significantly affecting glycolytic ATP production. Neither dapagliflozin nor empagliflozin reproduced these cellular and metabolic effects at any of the concentrations tested. SGLT2 and SGLT1 expression was low or undetectable in vascular tissues and cells, and neither sotagliflozin nor mizagliflozin reproduced the anti-calcifying effect of canagliflozin *in vivo* or *in vitro*.

**Conclusions:** Canagliflozin exhibits a selective anti-calcifying profile among the gliflozins tested, associated with changes in VSMC responses and cellular metabolism. These findings suggest that its vascular protective effects may involve mechanisms beyond canonical SGLT2 inhibition.

## Introduction

Chronic kidney disease (CKD) is associated with a substantial increase in cardiovascular morbidity and mortality.^1^ In addition to traditional cardiovascular risk factors such as hypertension, diabetes, and dyslipidemia, CKD-specific abnormalities, including mineral metabolism disorders, chronic inflammation, oxidative stress and uremic toxicity, contribute to vascular injury and accelerate vascular calcification.^2^ Vascular calcification is a strong predictor of cardiovascular events and mortality in patients with CKD, yet no therapy has been specifically established to prevent or reverse this process.^2–4^

CKD-associated vascular calcification is characterized by the pathological deposition of calcium-phosphate minerals within the vascular wall, particularly in the medial layer.^5^ This process increases arterial stiffness and pulse pressure, reduces vascular compliance, and contributes to left ventricular hypertrophy, heart failure, and cardiovascular mortality.^2,6,7^ Hyperphosphatemia is a major driver of CKD-associated vascular calcification by promoting calcium-phosphate crystal formation and providing a potent pro-calcifying stimulus to vascular smooth muscle cells (VSMCs).^8–10^ Rather than a passive precipitation phenomenon, vascular calcification is now recognized as an active and highly regulated cellular process in which VSMCs progressively lose contractile features and acquire osteogenic, inflammatory and stress-related phenotypes.^8,11^

Sodium-glucose cotransporter 2 inhibitors (SGLT2i), or gliflozins, were initially developed as glucose-lowering therapies for type 2 diabetes, through inhibition of renal glucose and sodium reabsorption in the proximal tubule.^12^ Large clinical trials subsequently demonstrated major cardiorenal benefits, including reduced CKD progression, heart failure hospitalization and cardiovascular mortality, in patients with and without diabetes.^13–18^ These benefits are only partly explained by glucose lowering and involve multiple mechanisms, including renal hemodynamic effects, reduced tubular workload, improved volume status, and modulation of inflammatory, oxidative and metabolic pathways.^19–21^ However, whether these established cardiorenal benefits extend to vascular calcification remains unclear. By slowing CKD progression, gliflozins may indirectly reduce the systemic pro-calcifying environment, while their reported effects on inflammation, oxidative stress, and cellular metabolism suggest potential direct actions on the vascular wall. Several experimental studies have reported anti-calcifying effects of individual gliflozins, including canagliflozin, dapagliflozin and empagliflozin, although findings have varied across experimental models and treatment conditions.^22–27^ This heterogeneity raises questions about the consistency of their vascular effects and the mechanisms involved.

An important unresolved question is whether these anti-calcifying effects are shared across gliflozins or depend on the individual compound. Although canagliflozin, dapagliflozin and empagliflozin preferentially inhibit SGLT2, they differ in their selectivity over the related sodium-glucose cotransporter SGLT1, which contributes to intestinal glucose absorption and renal glucose reabsorption. In particular, canagliflozin exhibits lower SGLT2/SGLT1 selectivity than dapagliflozin and empagliflozin.^28^ Beyond differences in transporter selectivity, distinct cellular effects have been reported for individual gliflozins, raising the possibility of compound-specific off-target mechanisms.^29–32^ Moreover, the involvement of canonical SGLT2 inhibition in the vascular effects of gliflozins remains uncertain, particularly given the inconsistent reports of SGLT2 expression in vascular cells.^33–35^

To address these questions, we performed a systematic comparison of canagliflozin, dapagliflozin and empagliflozin across complementary *in vivo*, *ex vivo* and *in vitro* models of vascular calcification, including human vascular tissues. We aimed to establish whether gliflozins share a common anti-calcifying effect or display molecule-specific activity, to characterize associated changes in VSMC proliferation, migration, glucose uptake, and cellular bioenergetics, and to examine the potential involvement of canonical SGLT2 and SGLT1 inhibition.

## Methods

### Animal experiments and ethical approval

Female DBA/2N mice (Charles River, France) were housed in a pathogen-free, temperature-controlled environment under a 12 h/12 h light/dark cycle, with free access to food and water. These mice have an inherent susceptibility to high-phosphate diet-triggered calcification.^36,37^ Upon arrival, mice were randomly assigned to cages (5 animals per cage), and cages were then randomly assigned to experimental groups. At 12 weeks of age, mice were fed for 2 weeks with either a standard diet (A04, Safe) or a phosphate-enriched diet (modified C1049, 2% phosphate, Altromin), according to a previously established model of high-phosphate diet-induced calcification.^38^

Two independent animal experiments were performed. In the first experiment, mice were assigned to five groups (10 animals per group): standard diet + vehicle (SD), high-phosphate diet + vehicle (HPD), HPD + canagliflozin, HPD + dapagliflozin or HPD + empagliflozin. Canagliflozin (MCE, HY-10451), dapagliflozin (MCE, HY-10450), empagliflozin (MCE, HY-15409) or vehicle was administered five days a week by oral gavage at 3mg/kg/day. Treatments were prepared at 0.3 mg/mL in a solution containing 5% DMSO, 40% PEG 300, 5% Tween-80, and 50% saline solution (vehicle). Treatments were administered in a blinded manner. One animal died in the HPD+dapa group. After 2 weeks, glomerular filtration rate (GFR) was measured in 7 animals per group. Urine samples were collected by bladder puncture and stored at −80°C. We were unable to obtain urine from one animal in the HPD group and one animal in the HPD+dapa group. At sacrifice, blood was collected from the vena cava into heparinized tubes (Microvette Hep-Lithium, Sarstedt) to prevent platelet aggregation. Blood was then centrifuged for 5 minutes at 1500 g, and plasma was collected and immediately frozen at −80 °C. One blood sample from the HPD group was excluded because of clot formation. The aortic arch, one renal pole, and the base of the heart were collected and immediately placed in 1 M HCl for assessment of calcification. Aortic and kidney samples were either snap-frozen in liquid nitrogen or fixed in paraformaldehyde solution and subsequently embedded in paraffin. Right femurs were dissected and placed in 70% ethanol for histomorphometric analyses.

In the second experiment, HPD-fed DBA/2N mice (6 animal per group) were treated with sotagliflozin (HY-15516), mizagliflozin (HY-17638) or vehicle at 3 mg/kg/day for 2 weeks using the same route of administration, dosing schedule and vehicle formulation. Urine was collected to assess glycosuria, and aortic arch was collected for calcium quantification. In addition, illustrative OsteoSense imaging of the whole aorta was performed. Mice received a retro-orbital injection of OsteoSense 750EX under isoflurane anesthesia 24 h before sacrifice. At sacrifice, the entire aorta was dissected from the aortic arch to the femoral artery bifurcation and placed in ice-cold PBS until imaging. Aortas were imaged in bright field and epifluorescence mode using an IVIS Spectrum In Vivo Imaging System (PerkinElmer, Waltham, MA, USA).

All experiments were performed in accordance with the National Institutes of Health guidelines for the care and use of laboratory animals and were approved by a local ethics committee (Centre Régional pour l’exploration fonctionnelle et de ressources expérimentales [CREFRE]/US006 Inserm, Toulouse, France; 23/1297/08/02 2023020717092839).

### Glomerular filtration rate

Mice were anesthetized with isoflurane, and a fluorescence measurement device (MediBeacon) was attached transdermally to previously depilated skin on the animals’ backs. An initial 1-minute measurement was performed to quantify baseline autofluorescence. Animals were then injected retro-orbitally with FITC-sinistrin (7 mg/100 g body weight) and individually placed in cages without access to water for 45 minutes. Fluorescence data were analyzed using MPD Lab software (v2.15, MediBeacon).

### Plasma and urine analyses

Plasma urea, calcium, and phosphate concentrations were determined using a biochemical analyzer (ABX Pentra C400, Horiba Medical). Complete blood counts were performed using a hematology analyzer (DF 56 Vet, Horiba Medical)). Plasma concentrations of parathyroid hormone (PTH) and fibroblast growth factor 23 (FGF23) were determined using the Mouse PTH 1-84 ELISA Kit (QuidelOrtho #60-2305) and MICROVUE™ Mouse/Rat FGF-23 ELISA (C-Term) (QuidelOrtho #60-300), respectively, according to the manufacturer’s instructions. Blood glucose levels were measured from tail blood under isoflurane anesthesia using an Accu-chek glucometer (07400942016) after 6h of fasting.

Urinary glucose was quantified using the Glucose GOD FS test (Eurobio, 1 2500 99 10 923), according to the manufacturer’s instructions. Urine samples were incubated with the detection reagent for 20 minutes at 25°C, after which absorbance at 500 nm was measured using a SpectroStar Omega plate reader (BMG Labtech). Urinary albumin was measured using the AlbuWell kit (Ethos Biosciences, 1011). Urinary phosphate concentration was determined using the QuantiChrom Phosphate Assay kit (BioAssay Systems, DIPI-500) according to the manufacturer’s instructions. Urinary creatinine was measured using the Jaffe colorimetric method with the Creatinine Assay kit (BioAssay Systems, DICT-500) for normalization.

### Culture of primary human and murine aortic smooth muscle cells

Primary human aortic smooth muscle cells (hASMCs, Lonza, CC-2571) and mouse VSMCs (MOVAS, ATCC-CRL 2797) were maintained at 37°C in a humidified atmosphere containing 5% CO₂, in SMGM™ BulletKit medium (Lonza, CC-3182) or DMEM GlutaMAX (Gibco 61965-026), respectively, supplemented with 10% fetal bovine serum (FBS), 100 U/mL penicillin, and 100 µg/mL streptomycin until confluence. For experiments, cells were split and cultured in DMEM GlutaMAX supplemented with 2.5% FBS, 100 U/mL penicillin, and 100 µg/mL streptomycin.

To induce calcification, cells were treated with 2.6 mM Na₂HPO₄ / 25 mM HEPES for 5 days (calcium assay and OsteoSense staining). As indicated, hASMCs and MOVAS cells were also treated with 1.25-10 µM canagliflozin (MCE HY-10451), dapagliflozin (MCE HY-10450), empagliflozin (MCE HY-15409), mizagliflozin (MCE HY17638) or sotagliflozin (NB-64-23145) for 5 days. Equivalent amounts of vehicle were used as control. Fresh medium containing the compounds was replaced every two days. hASMCs from male and female donors (tissue acquisition #26646, #35724), as well as MOVAS cells, were used between passages 3 and 7 depending on availability. All experiments were performed using three to four independent cell passages, with at least 4 biological replicates per group.

### Culture of mouse and rat aortic rings

Thoracic aortas were dissected from 12-week-old male C57BL/6 mice, diabetic db/db mice and their non-diabetic db/+ controls (Janvier Labs), and 6-week-old Wistar Han male rats (Charles River Laboratories). Connective tissue was carefully removed, and the aortas were cut into 2-3 mm-long rings. Rings from the same animal were randomly allocated among the different experimental conditions. Rings were cultured in DMEM supplemented with 2.5% FBS, 100 U/mL penicillin, and 100 µg/mL streptomycin at 37°C in a humidified atmosphere containing 5% CO₂.

To induce calcification, rings were stimulated with 1.8 mM Na₂HPO₄ / 25 mM HEPES for 7 days (mice, calcium assay) or 14 days (rats, von Kossa staining). Where indicated, aortic rings were also treated with 1.25 and 5 µM canagliflozin (MCE HY-10451), dapagliflozin (MCE HY-10450), or empagliflozin (MCE HY-15409). Equivalent amounts of vehicle were used as control. Fresh medium was replaced every two days.

### Culture of human vascular explants

Six patients with stage 5 CKD (GFR <15 mL/min) eligible for living-donor kidney transplantation were enrolled in the Department of Nephrology at Karolinska University Hospital, Stockholm, Sweden. The study was approved by the Regional Ethical Review Board in Stockholm (approval numbers: Dnr 2015/1115-31, amendment 2017/1799-32; Dnr 2008/1748-31/2, amendment 2015/1789/32; and Dnr 2011/668-31, amendments 2014/968-32 and 2019-02912). Written informed consent was obtained from all participants prior to tissue collection. All procedures were conducted in accordance with the principles of the Declaration of Helsinki. During transplantation, fragments of the iliac and epigastric arteries were collected and cut into 2-3 mm rings. Subcutaneous adipose tissue was also collected from the incision site, from which resistance arteries (diameter ≈150–400 µm) were dissected. In addition, aortic explants were obtained from deceased organ donors during kidney transplantation procedures in the Department of Nephrology, Dialysis and Transplantation at Rangueil University Hospital, Toulouse. All donors were included in the Nephrogene Cohort (RC31/21/0154), which authorizes the use of biological samples for research purposes. Written informed consent was obtained in each case. Cross-sectional pieces measuring 5 × 5 mm were prepared. Aortic patches from donors with a history of atherosclerosis or diabetes were excluded from the study. Rings/sections from the different vascular territories of each patient were allocated among the different experimental conditions. The number of replicates per condition varied between patients depending on the availability and size of the available samples.

At collection, vessels were immediately transferred into PBS at 4 °C, then retrieved and cultured in DMEM supplemented with 2.5% FBS, 100 U/mL penicillin, and 100 µg/mL streptomycin at 37°C in a humidified atmosphere containing 5% CO₂. To induce calcification, rings were stimulated with 1.8 mM Na₂HPO₄ / 25 mM HEPES for 7 days. Where indicated, aortic rings were also treated with 1.25 and 5 µM canagliflozin (MCE HY-10451), dapagliflozin (MCE HY-10450), or empagliflozin (MCE HY-15409). Equivalent amounts of vehicle were used as control. Fresh medium was replaced every two days.

### Quantification of calcium deposits

Cells and tissues were decalcified overnight in 1 M HCl at 4°C. Calcium concentration in the supernatant was determined using the QuantiChrom Calcium Assay kit (BioAssay Systems, DICA-500) according to the manufacturer’s instructions. Cells were subsequently lysed with 1 M NaOH/0.1% SDS, and protein concentration was measured using the Pierce BCA kit (Thermo Scientific) for normalization. Tissues (vessels, kidneys, and hearts) were weighed. Results are presented after normalization to total protein concentration or tissue weight.

### Detection of calcification using OsteoSense

For detection of calcification in MOVAS cells, after 5 days of treatment, the OsteoSense™ 680 probe (NEV10020EX) was added directly to the culture medium at a final concentration of 1 pmol/mL. After 24 h of incubation, cells were washed twice with PBS, and the signal was acquired using the IncuCyte® SX5 real-time imaging system (Sartorius, Controller Version 2025B).

### Cell proliferation and migration monitoring

Cell proliferation and migration was monitored by real-time imaging using the IncuCyte® SX5 real-time imaging system (Sartorius). For proliferation assay, MOVAS cells were seeded in 96-well plates at low confluence (100 cells per well) and then cultured under the indicated experimental conditions. Plates were placed in the IncuCyte®, maintained at 37 °C under 5% CO₂, and phase-contrast images were automatically acquired at regular 6-h intervals for 5 days. Proliferation was assessed by automated cell counting using IncuCyte® software.

For cell migration scratch wound analysis, MOVAS cells were seeded at a density of 25,000 cells per well in Incucyte® ImageLock 96-well plates (Sartorius BA-04856) to achieve 100% confluence after 24 h. The following day, uniform scratch wounds were generated using the Incucyte® 96-Well Woundmaker Tool (Sartorius 4563). After wounding, the culture medium was replaced with indicated experimental conditions. Wound closure was monitored for 48 h using the Incucyte® Live-Cell Analysis System, with phase-contrast images acquired every 2 h. Cell migration was quantified by measuring relative wound density (RWD) over time using the Incucyte® Scratch Wound analysis software module (Sartorius, Cat. No. 9600-0012).

### Cell death monitoring

For detection of apoptotic cells, MOVAS cells were stained with annexin V-iFluor® 488 and Propidium Iodide (AAT Bioquest 22824, 100× stock solutions), diluted in DMEM containing 1% FBS, for 30 minutes after 2 and 7 days of treatment. The green and orange fluorescent signal was quantified using the IncuCyte® system. Positive staining was normalized to the total number of cells per condition.

### Seahorse XF Real-Time ATP rate assay

Extracellular ATP production was assessed using the Seahorse XF Real-Time ATP Rate Assay Kit (Agilent Technologies, Cat. No. 103592-100). MOVAS cells were seeded at a density of 5,000 cells per well in Seahorse XF96 cell culture microplates (Agilent Technologies). After 24 h, cells were exposed to the different experimental conditions for 48 h. Following treatment, the culture medium was replaced with Seahorse XF DMEM Medium (Agilent Technologies) supplemented with 10 mM glucose, 1 mM pyruvate, and 2 mM glutamine. Mitochondrial and glycolytic ATP production rates were then assessed using a Seahorse XF Pro 96 Analyzer (Agilent Technologies), according to the manufacturer’s instructions. Briefly, oxygen consumption rate (OCR) and extracellular acidification rate (ECAR) were measured at baseline and following sequential injections of oligomycin (1.5 µM), an ATP synthase inhibitor, and rotenone/antimycin A (0.5 µM), inhibitors of mitochondrial complexes I and III, respectively. Mitochondrial ATP production rate (mitoATP), glycolytic ATP production rate (glycoATP), and total ATP production rate were calculated using the Seahorse XF Real-Time ATP Rate Assay analysis software (Agilent Technologies). ATP production rates were normalized to cell number, as quantified using the Incucyte® Live-Cell Analysis System (Sartorius).

### Glucose uptake assay

Glucose uptake was assessed using radiolabeled 2-deoxy-D-glucose ([³H]-2-DG). MOVAS cells were seeded at a density of 100.000 cells per well in 12-well plates. The following day, cells were treated with canagliflozin, dapagliflozin, or empagliflozin (1.25, 5, or 10 µM) for 1 h in DMEM 4.5 g/L glucose (Gibco 61965-026) supplemented with 2.5% (FBS). Cells were then washed with PBS and incubated for an additional 1 h in serum- and glucose-free DMEM (Gibco 11966-025), while maintaining the respective treatments. Following glucose deprivation, the medium was replaced with 450 µL of glucose-free DMEM without SGLT2 inhibitors. Glucose uptake was initiated by adding 50 µL of a mixture containing unlabeled 2-deoxy-D-glucose (2-DG) and [1,2-³H(N)]-2-deoxy-D-glucose (PerkinElmer, Cat. No. NET549005MC), resulting in a final concentration of 137.5 µM unlabeled 2-DG and 0.2 µCi of [³H]-2-DG per well. After 30 min of incubation, cells were washed and then lysed with 0.05 N NaOH. Intracellular radioactivity was quantified using a Tri-Carb liquid scintillation counter (PerkinElmer). Glucose uptake rates were calculated based on the effective specific activity of the tracer mixture and normalized to total protein content determined using the Pierce BCA kit (Thermo Scientific) for normalization. Results were expressed as nmol of internalized 2-DG/min/mg of protein.

### Immunohistochemical and histological analyses

Four-micrometer-thick mouse kidney sections were deparaffinized and rehydrated. After blocking endogenous peroxidases (Dual Endogenous Enzyme Block, Dako, 10 minutes), sections were incubated with the primary antibody (Table 1) overnight at 4 °C, washed with TBS containing 0.1% Tween 20, and incubated for 1 h with the secondary antibody (Table 1). Detection was performed using a 3,3′-diaminobenzidine (DAB) chromogenic substrate solution (Liquid DAB + Substrate Chromogen System, Dako), followed by hematoxylin counterstaining. In addition, for detection of calcification, kidney sections were stained with 2% Alizarin Red (Merck) (pH 4.2) for 45 minutes. Finally, slides were dehydrated and mounted under coverslips. Sections were scanned using a NanoZoomer 2.0 RS (Hamamatsu Photonics SARL, Massy, France), and positive staining was quantified using ImageJ1 Fiji software (https://imagej.net/downloads). The number of stained pixels was then expressed relative to the total number of pixels covering the section area.

**Table 1.** Primary and secondary antibodies used for IHC and IF.

| **Primary antibodies** | **Secondary antibodies** |
|---|---|
| F4/80 (MF 48000, Invitrogen), 1/100 | Anti-rat HRP (414311F, Histofine), 1/2 |
| Collagen 3 (ab59436, Abcam), 1/200 | Anti-Rabbit HRP (414341F, Histofine), 1/2 |
| Sglt2 (24654-1-AP, Proteintech), 1/200 | Goat anti-Rabbit IgG (H+L), Alexa Fluor 488 (ThermoFisher A11008), 1/1000 |
| Sglt1 (07-1417, Sigma), 1/200 |  |
| Alexa Fluor 647anti-α-Smooth Muscle Actin | Not applicable |

Femurs were dehydrated in alcohol, treated with xylene, and then embedded in a mixture consisting of 75% methyl methacrylate, 25% dibutyl phthalate, and 2.5% (w/v) benzoyl peroxide. Histomorphometric parameters were assessed on 5-µm undecalcified sections of the distal end, stained using a modified Masson-Goldner trichrome technique. Briefly, sections were fixed with 50% ethanol under pressure, rehydrated, and then stained with a hematoxylin–ferric chloride mixture (1:1), followed by differentiation with 1% hydrochloric acid (HCl) and bluing with lithium carbonate (LiCO₃). After rinsing with water, sections were stained with Goldner trichrome for 20 minutes and then rinsed with 1% acetic acid. Samples were subsequently stained with safranin for five minutes, dehydrated in ethanol, and mounted. Green-stained areas were considered mineralized bone, whereas red-stained areas at least 1.5 µm thick were identified as osteoid. Histomorphometric parameters were analyzed in trabecular bone within the secondary spongiosa (at a distance of 0.25 mm from the endocortical bone and growth plate), at 40× and 100× magnification, using OsteoMeasure software (OsteoMetrics, USA), and derived indices were calculated according to standard methods. Dynamic bone parameters were determined by measuring single and double fluorescent calcein labeling lines on serial 10-µm undecalcified sections.

Rat aortic rings were fixed in 4% formaldehyde, embedded in paraffin, and cut into 5-µm sections. After deparaffinization and rehydratation, von Kossa staining (silver nitrate and nuclear red) was performed and sections were photographed (NIKON TM300 microscope equipped with a DXM1200 digital imaging system) at 40-fold magnification. The positively stained area was quantified within the media and expressed as a percentage of the total area.

### Immunofluorescence staining

For detection of SGLT2 and SGLT1, aortic and kidney samples were embedded in OCT medium at −80°C and then cut into 5-µm-thick cryosections, while MOVAS cells were cultured on coverslips. After fixation with 4% paraformaldehyde, blocking of nonspecific binding sites, and permeabilization (0.05% Triton X-100), sections and cells were incubated overnight at 4°C with primary antibodies directed against SGLT1 or SGLT2 (Table 1). For aortas, co-staining was performed using a directly conjugated anti-α-SMA antibody (Alexa Fluor 647) to label the media. After washing, samples were incubated with an Alexa Fluor® 488-conjugated secondary antibody, and nuclei were counterstained with DAPI. Staining was acquired using a Zeiss Cell Observer video microscope (Zeiss).

### Gene expression analysis

Total RNA was extracted from kidney cortex and duodenum using the RNeasy Mini Kit (Qiagen), according to the manufacturer’s instructions. Total RNA from murine thoracic aorta, MOVAS cells, hASMC, and SV40T-immortalized human mesenteric artery-derived VSMC^39^ was extracted using TRIzol™ reagent (ThermoFisher Scientific) according to the manufacturer’s protocol. Samples were homogenized in TRIzol, followed by phenol-chloroform extraction. After centrifugation, the aqueous phase was collected, and RNA was precipitated with isopropanol, washed with 75% ethanol, and resuspended in RNase-free water. RNA concentration was measured using a NanoDrop ND-2000 spectrophotometer (Thermo Fisher Scientific, Waltham, MA, USA). Total RNA was reverse-transcribed using the Maxima First Strand cDNA Synthesis Kit (ThermoFisher Scientific). RT-qPCR was performed using the QuantStudio Design & Analysis software system v1.5.1 with ONEGreen FAST qPCR Premix (Ozyme), according to the manufacturer’s instructions. The primer sequences used in this study are listed in Table 2. For mouse and human SGLT1/SGLT2 expression analysis, RT-qPCR was performed using TaqMan Gene Expression Assays (Life Technologies/Thermo Fisher Scientific) with TaqMan Universal PCR Master Mix II (catalog no. 4440040). The following assays were used: human *SLC5A1* / SGLT1, Hs01573793_m1; mouse *Slc5a1* / SGLT1, Mm00451203_m1; human *SLC5A2* / SGLT2, Hs00894642_m1; and mouse *Slc5a2* / SGLT2, Mm00453831_m1. Relative gene expression was calculated using the 2^-ΔΔCt method, with Gapdh and Hprt as reference genes.

**Table 2.**
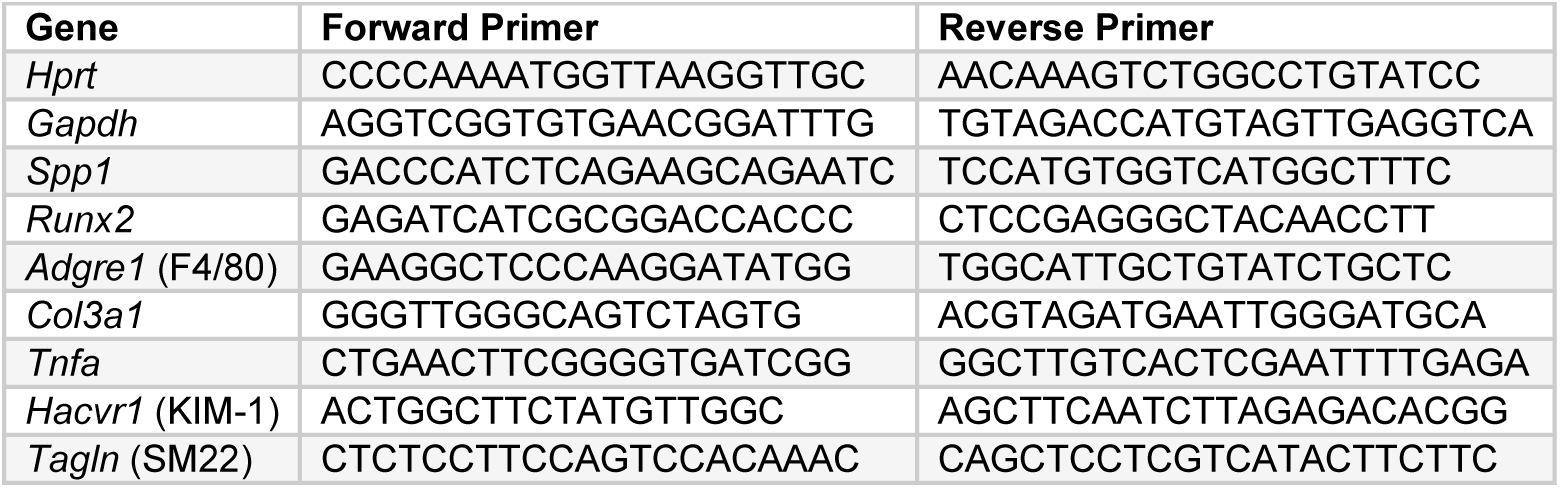
Primer sequences used (species: mouse)

### RNA sequencing

Total RNA was extracted from mouse aortas using TRIzol/chloroform, with three biological replicates per experimental group. RNA concentration and purity were assessed using a NanoDrop ND-2000 spectrophotometer (Thermo Fisher Scientific, Waltham, MA, USA), and RNA integrity was evaluated using a Fragment Analyzer with the RNA Standard Sensitivity Kit (Agilent Technologies, Santa Clara, CA, USA).

Strand-specific RNA-seq libraries were prepared from 500 ng of total RNA using the TruSeq Stranded mRNA Library Prep Kit (Illumina, San Diego, CA, USA). Following poly(A) selection, RNA fragmentation, cDNA synthesis, and adapter ligation, libraries were amplified by 14 PCR cycles. Library quality was assessed using a Fragment Analyzer with the HS NGS Kit (Agilent Technologies), and concentrations were determined using the Qubit dsDNA HS Assay Kit on a Qubit 3.0 fluorometer (Thermo Fisher Scientific). Libraries were pooled at equimolar concentrations and subjected to five additional amplification cycles using the KAPA HiFi HotStart Library Amplification Kit (Roche, Basel, Switzerland) to ensure compatibility with AVITI circularization. Paired-end sequencing (2 × 150 bp) was performed at the GeT-PlaGe facility (INRAE, Toulouse, France) using an AVITI sequencer (Element Biosciences, San Diego, CA, USA).

### Western blot

Total proteins were extracted from kidney and aortic tissues of wild-type (WT) and SGLT2 knockout (KO) mice kindly provided by the lab of Prof. Coert J. Zuurbier (Amsterdam UMC, The Netherlands), whose previous studies employed this SGLT2-deficient mouse model^40,41^, as well as from MOVAS cells. Equal amounts of protein (30 µg) were separated by SDS-polyacrylamide gel electrophoresis (SDS-PAGE) using Stain-Free precast gels (Bio-Rad) and subsequently transferred onto nitrocellulose membranes (Amersham) using a Trans-Blot transfer system (Bio-Rad). Membranes were blocked for 1 h at room temperature in Tris-buffered saline containing 0.1% Tween-20 (TBST) and 5% non-fat dry milk, followed by overnight incubation at 4°C with primary antibodies against SGLT2 (proteintech 24654-1-AP, 1/1000) or GAPDH (Cell signaling 2118, 1/1000), diluted in TBST containing 5% non-fat dry milk. Membranes were then incubated with a horseradish peroxidase (HRP)-conjugated anti-rabbit secondary antibody (cell signaling #7074, 1/10000) for 1 h at room temperature. Immunoreactive bands were detected using enhanced chemiluminescence (ECL) reagents (Amersham), and images were acquired using a ChemiDoc imaging system (Bio-Rad). Densitometric analysis was performed using Image Lab software (Bio-Rad).

### Statistical analyses

Statistical analyses of experimental data were performed using GraphPad Prism 11.0 software (GraphPad Software, San Diego, California, USA, www.graphpad.com). All data are presented as mean ± standard error of the mean. Statistical tests were selected based on assessment of data distribution and homoscedasticity. Data normality was assessed using the Shapiro-Wilk test. Homogeneity of variances was evaluated using an F-test for two-group comparisons and the Brown-Forsythe test for comparisons among three or more groups. For comparisons between two independent groups, normally distributed data were analyzed using an unpaired Student’s t-test when variances were homogeneous or Welch’s t-test when variances were unequal. Non-normally distributed data were analyzed using the Mann–Whitney U test. For comparisons among three or more groups, normally distributed data were analyzed using one-way ANOVA followed by Dunnett’s multiple comparisons test when variances were homogeneous, or Brown–Forsythe and Welch ANOVA followed by Dunnett’s T3 multiple comparisons test when variances were unequal. Non-normally distributed data were analyzed using the Kruskal-Wallis test followed by Dunn’s multiple comparisons test. For the simultaneous assessment of two factors, two-way ANOVA was performed followed by Dunnett’s correction for multiple comparisons. In all cases, p < 0.05 was considered statistically significant.

RNA-sequencing analysis was performed in R (v4.5.2), using the edgeR (v4.8.2) and limma (v3.66.0) packages. Lowly expressed genes were removed using filterByExpr, accounting for library sizes and group structure. Library normalization factors were subsequently calculated using the trimmed mean of M-values (TMM) method. Gene counts were transformed to log2 counts per million, and the mean-variance relationship was estimated using the voom method. Linear models were fitted using limma, with experimental groups represented in the design matrix. Predefined comparisons between conditions were specified using makeContrasts, and empirical Bayes moderation was applied to obtain moderated t-statistics and associated p-values. Differentially expressed genes were identified using an adjusted p-value (Benjamini–Hochberg method) below 0.05. Results were represented as volcano plots (log2 fold-change vs. −log10 p-value) and heatmaps of differentially expressed genes based on z-score-standardized expression values.

Finally, gene set enrichment analysis (GSEA) was performed using a preranked approach, with genes ranked according to the moderated t-statistics obtained from the differential expression analysis. Analyses were run without gene-symbol collapsing, using the mouse Hallmark gene sets from the Molecular Signatures Database (MSigDB) Hallmark collection (mh.all.v2026.1.Mm.symbols.gmt). Pathway enrichment was assessed using the normalized enrichment score (NES) and FDR q-value.

## Results

### High-phosphate diet induces renal dysfunction, nephrocalcinosis and CKD-MBD features in DBA/2N mice

To compare the effects of different gliflozins on phosphate-driven ectopic calcification, female DBA/2N mice were fed either a standard diet (SD) or a high-phosphate diet (HPD) for 2 weeks and treated with canagliflozin, dapagliflozin or empagliflozin at 3 mg/kg/day (**Fig. 1A**). This dose was selected based on pharmacodynamic validation showing that all three compounds decreased blood glucose and increased urinary glucose excretion in diabetic db/db mice (**Supplementary Fig. S1**), and increased urinary glucose excretion in DBA/2N mice in the experimental setting used in this study (**Supplementary Fig. S1** and **Fig. 1B**).

**Figure 1.**
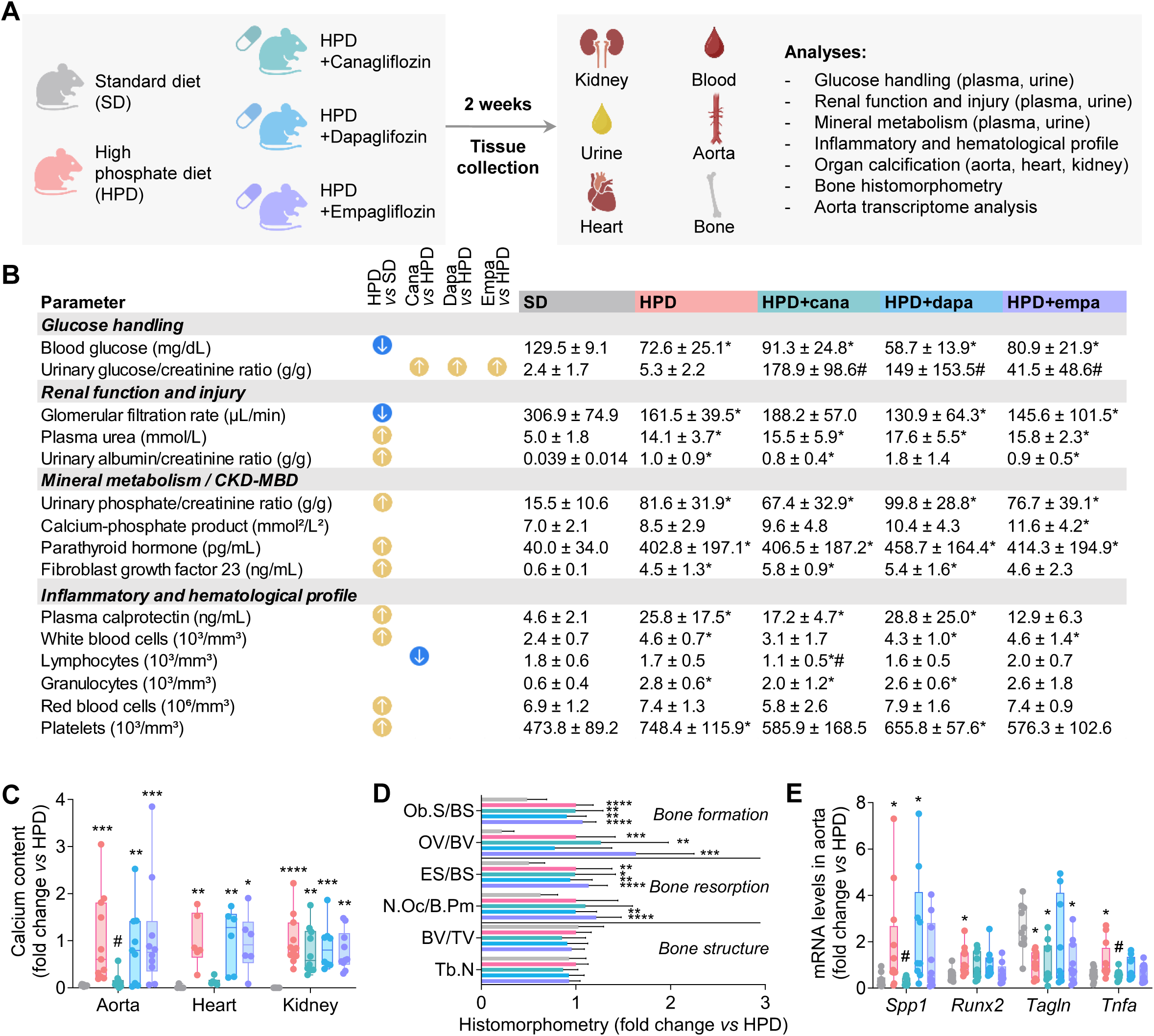
Canagliflozin attenuates high-phosphate diet-induced vascular calcification. **(A)** Experimental design. Female DBA/2N mice were fed either a standard diet (SD) or a high-phosphate diet (HPD) and were treated with canagliflozin, dapagliflozin or empagliflozin for 2 weeks. Blood, urine and tissues relevant to CKD-associated vascular calcification were collected for phenotypic, histological and molecular analyses. The panel was created in part using BioRender.com. **(B)** Summary table of glucose handling, renal function and injury, mineral metabolism/CKD-MBD, and inflammatory and hematological parameters across experimental groups. **(C)** Calcium content in the aortic arch, heart and kidney across experimental groups. **(D)** Bone histomorphometry assessing bone formation, bone resorption and trabecular bone structure, including osteoblast surface per bone surface (Ob.S/BS), osteoid volume per bone volume (OV/BV), eroded surface per bone surface (ES/BS), osteoclast number per bone perimeter (N.Oc/B.Pm), bone volume per trabecular volume (BV/TV) and trabecular number (Tb.N). Data are shown as mean ± SD. **(E)** Aortic mRNA expression of osteogenic, contractile and inflammatory markers measured by RT-qPCR: *Spp1* (secreted phosphoprotein 1/osteopontin), *Runx2* (runt-related transcription factor 2), *Tagln* (transgelin/SM22α), and *Tnfa* (TNFα). Data are displayed in mean ± SD. *P < 0.05, **P < 0.01, ***P < 0.001 and ****P < 0.0001 versus SD; #P < 0.05 versus HPD.

Renal function and injury, mineral and bone disorder (MBD), and inflammatory and hematological parameters across experimental groups are summarized in **Fig. 1B**. Compared with SD mice, HPD animals developed renal dysfunction and injury, with reduced glomerular filtration rate, increased plasma urea and increased urinary albumin-to-creatinine ratio. HPD also increased urinary phosphate excretion and plasma levels of intact parathyroid hormone (PTH 1-84) and C-terminal fibroblast growth factor 23 (FGF23), consistent with CKD-MBD-like abnormalities. In parallel, HPD modified the inflammatory and hematological profile, as shown by increased plasma calprotectin, white blood cells, granulocytes and platelets.

Consistent with this phenotype, HPD increased renal mRNA expression of *Havcr1* encoding KIM-1, *Adgre1* encoding F4/80, and *Col3a1*, together with increased renal F4/80 and type III collagen immunostaining (**Supplementary Fig. S2A-D**). Conversely, renal mRNA expression of *Slc5a2* encoding SGLT2, and SGLT2 immunostaining were reduced in HPD mice compared with SD mice (**Supplementary Fig. S2A-D**). HPD also induced marked renal mineral deposition, as shown by increased Alizarin Red staining, consistent with nephrocalcinosis (**Supplementary Fig. S2C, D**). Gliflozin treatment did not significantly modify renal function, renal injury, phosphate handling, CKD-MBD parameters, renal inflammatory/fibrotic markers or renal mineral deposition (**Fig. 1B** and **Supplementary Fig. S2**). Canagliflozin showed a trend toward lower systemic inflammatory and hematological parameters, including a significant reduction in circulating lymphocytes compared with HPD (**Fig. 1B**).

### Canagliflozin selectively attenuates HPD-induced vascular calcification

In this model, HPD induced marked ectopic calcification deposition in the aorta, heart and kidney (**Fig. 1C**). Among the three gliflozins, canagliflozin was the only treatment that significantly reduced aortic calcification. Canagliflozin also tended to reduce cardiac calcification, although this effect did not reach statistical significance. Dapagliflozin and empagliflozin did not significantly reduce tissue calcium deposition (**Fig. 1C**).

Bone histomorphometry was then performed to assess physiological mineralized tissue. HPD increased both bone formation and bone resorption, without major changes in global bone structure (**Fig. 1D** and **Supplementary Fig. S2E**). Gliflozin treatment did not significantly alter these bone parameters.

At the vascular molecular level, HPD induced a pro-calcifying and inflammatory transcriptional profile in the aorta, characterized by increased expression of *Spp1* (secreted phosphoprotein 1/osteopontin), *Runx2* (runt-related transcription factor 2) and *Tnfa* (TNFα), together with reduced expression of the contractile marker *Tagln* (transgelin/SM22α) (**Fig. 1E**). Canagliflozin significantly reduced aortic *Spp1* and *Tnfa* expression, but did not modify *Runx2* or *Tagln* expression. Neither dapagliflozin or empagliflozin affected these markers. Overall, canagliflozin was the only gliflozin that significantly attenuated aortic calcification in HPD-fed mice.

### Canagliflozin partially reverses HPD-induced aortic transcriptional alterations

To further characterize the vascular transcriptional response, RNA-seq was performed on aortic tissue. HPD profoundly remodeled the aortic transcriptome compared with SD, with 1,986 differentially expressed genes. Canagliflozin also induced a broad transcriptional response, with 1,090 differentially expressed genes compared with HPD alone. In contrast, dapagliflozin and empagliflozin had more limited effects, with 112 and 419 differentially expressed genes, respectively (**Fig. 2A**). Heatmap analysis showed that the top genes modified by canagliflozin versus HPD (selected using FDR q-value < 0.05 and absolute log₂ fold change ≥ 1.5) partially reversed the HPD-induced transcriptional pattern, whereas dapagliflozin and empagliflozin had limited effects on these genes. Conversely, the top genes modified by HPD versus SD were not globally normalized by gliflozin treatment (**Fig. 2B**).

**Figure 2.**
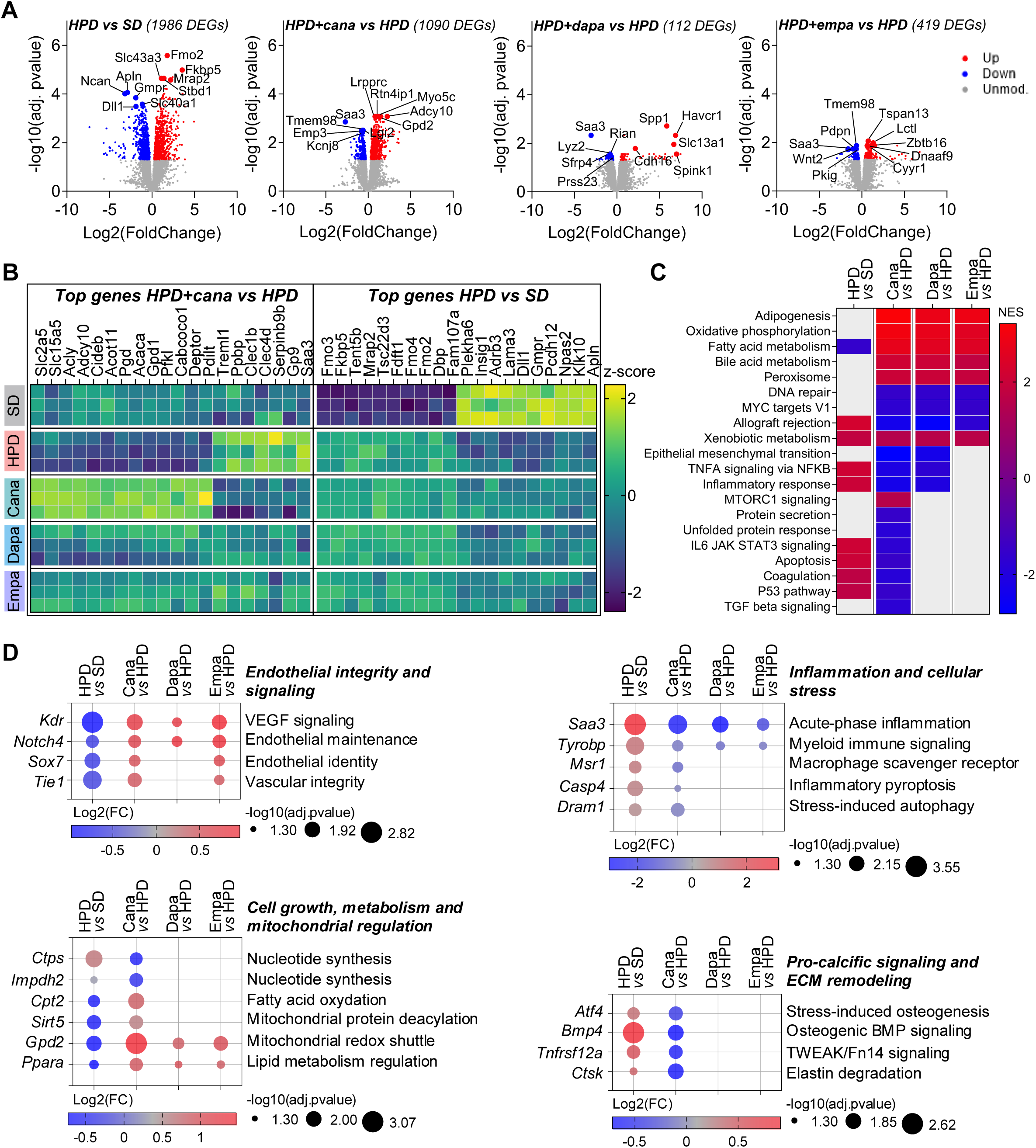
Canagliflozin partially reverses high-phosphate diet-induced aortic transcriptional changes. Aortic RNA sequencing (RNA-seq) was performed to characterize transcriptional alterations induced by HPD and their modulation by canagliflozin, dapagliflozin, and empagliflozin. **(A)** Volcano plots showing differentially expressed genes (DEGs) in HPD versus SD, HPD + canagliflozin versus HPD, HPD + dapagliflozin versus HPD, and HPD + empagliflozin versus HPD. **(B)** Heatmap showing the top 20 differentially expressed genes in HPD + canagliflozin versus HPD and the top 20 differentially expressed genes in HPD versus SD, selected using FDR q-value < 0.05 and absolute log₂ fold change ≥ 1.5. **(C)** Heatmap of the top enriched Hallmark pathways identified by preranked GSEA in HPD + canagliflozin versus HPD, and their corresponding enrichment patterns in HPD versus SD, HPD + dapagliflozin versus HPD, and HPD + empagliflozin versus HPD. Pathways were selected using FDR q-value < 0.10 and absolute normalized enrichment score (NES) ≥ 1.5. **(D)** Dot plots showing log2 fold changes and -log10(adj.pvalue) of selected genes significantly altered by HPD and significantly regulated in the opposite direction by canagliflozin. Fold changes induced by dapagliflozin and empagliflozin are also presented when statistically significant.

Pathway-level analysis by preranked GSEA further supported this selective transcriptional response (**Fig. 2C**). Several metabolic pathways, including adipogenesis, oxidative phosphorylation, and fatty acid metabolism, were modulated by multiple gliflozins. Canagliflozin additionally affected pathways overlapping with HPD-driven alterations, including inflammatory and apoptosis-related programs.

To explore potential mechanisms, we examined HPD-responsive genes significantly reversed by canagliflozin (**Fig. 2D**). HPD reduced the expression of genes associated with endothelial integrity and signaling (*Kdr*, *Notch4*, *Sox7*, and *Tie1*) and increased inflammatory markers (*Saa3*, *Tyrobp*). These changes were reversed by canagliflozin, with similar responses also observed following dapagliflozin and empagliflozin treatment. In contrast, canagliflozin significantly reversed changes in genes associated with pro-calcific signaling and matrix remodeling (*Atf4*, *Bmp4*, *Tnfrsf12a*, *Ctsk*), cell growth (*Ctps*, *Impdh2*), and mitochondrial metabolism (*Cpt2*, *Sirt5*), without significant effects of the other gliflozins on these genes.

Together, these findings indicate that canagliflozin partially reverses HPD-induced aortic transcriptional alterations. While some endothelial and inflammatory responses were shared among gliflozins, canagliflozin also modulated genes associated with pro-calcific signaling, cell growth, and metabolism, suggesting additional mechanisms underlying its vascular protection.

### Canagliflozin directly reduces vascular calcification ex vivo and in vitro

To determine whether this protection involves direct effects on the vascular wall, we next investigated the anti-calcifying activity of canagliflozin in isolated vascular tissues and cultured vascular smooth muscle cells. Gliflozins were therefore tested in complementary *ex vivo* and *in vitro* calcification models, including C57BL/6 and diabetic mouse aortic rings, rat aortic rings, human vascular explants from different vascular beds, primary human aortic smooth muscle cells (hASMCs) and MOVAS cells (**Fig. 3A**).

**Figure 3.**
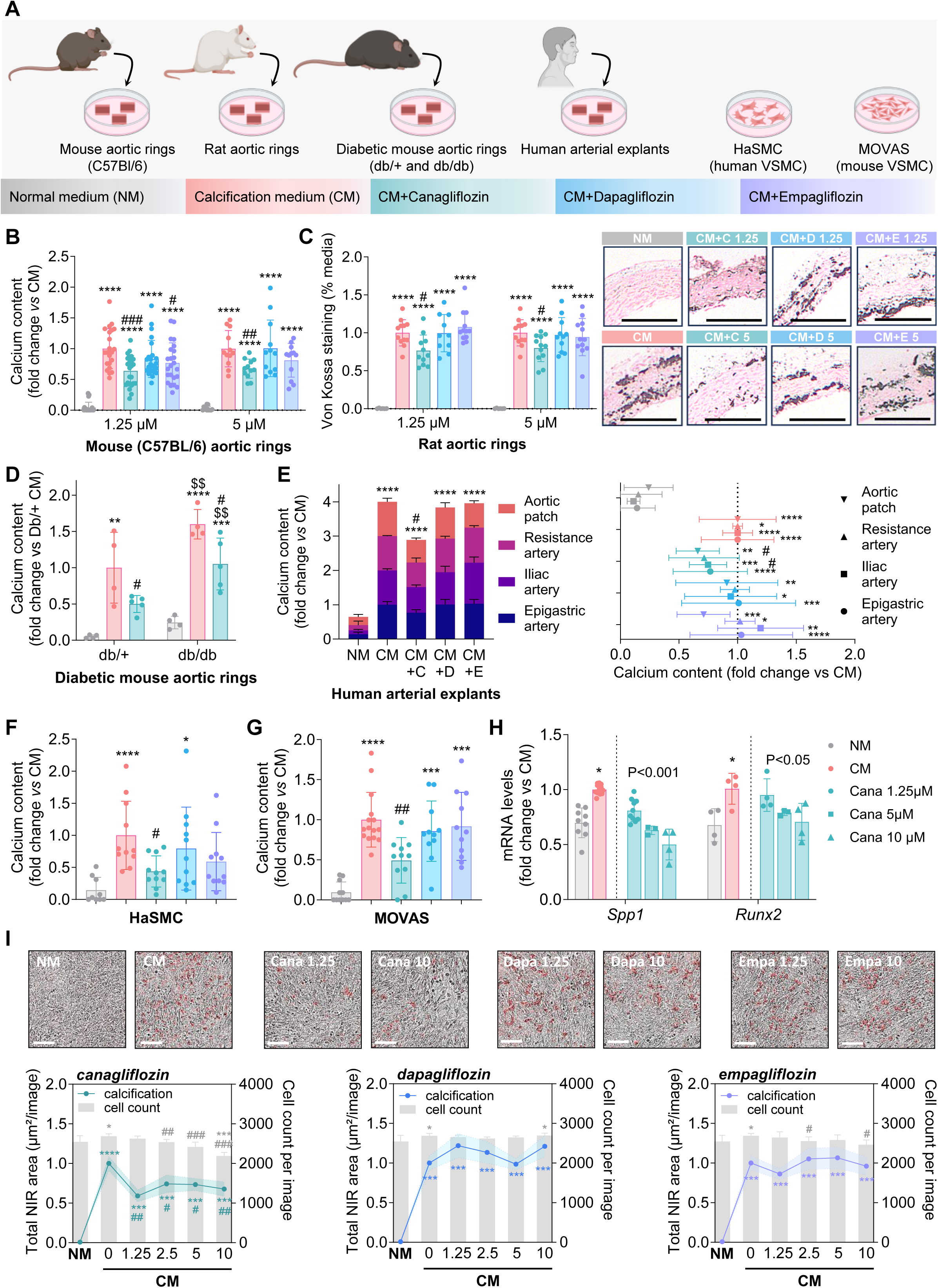
Ex vivo and in vitro effects of gliflozins on vascular calcification. **(A)** Overview of the experimental models used to assess the anti-calcifying effects of gliflozins, including control and diabetic mouse aortic rings, rat aortic rings, human arterial explants, primary human aortic smooth muscle cells (haSMCs) and MOVAS cells. The panel was created in part using BioRender.com. **(B)** Calcium content in C57BL/6 mouse aortic rings cultured under calcifying conditions and treated with canagliflozin, dapagliflozin or empagliflozin (1.25 or 5 µM). **(C)** Calcification analysis in rat aortic rings treated with canagliflozin, dapagliflozin or empagliflozin (1.25 or 5 µM), assessed by von Kossa staining of the vascular media. Quantification and representative images are shown, scale bar = 200µm. **(D)** Calcium content in aortic rings from non-diabetic db/+ and diabetic db/db mice cultured under calcifying conditions and treated with canagliflozin (1.25 µM). **(E)** Calcium content in human vascular explants, including aortic patches, resistance arteries, iliac arteries and epigastric arteries, treated with canagliflozin (CM+C), dapagliflozin (CM+D) or empagliflozin (CM+E) (5 µM). **(F)** Calcium content in primary hASMCs cultured under calcifying conditions and treated with canagliflozin, dapagliflozin or empagliflozin (1.25 µM). **(G)** Calcium content in MOVAS cells cultured under calcifying conditions and treated with canagliflozin, dapagliflozin or empagliflozin (1.25 µM)**. (H)** Relative mRNA expression of *Spp1* and *Runx2* in MOVAS cells cultured in calcifying conditions and treated with canagliflozin (1.25, 5 or 10 µM), assessed by RT-qPCR. **(I)** Dose-response analysis of MOVAS calcification under calcifying conditions after treatment with canagliflozin, dapagliflozin or empagliflozin (1.25, 2.5, 5 or 10 µM), assessed by OsteoSense staining, with quantification and representative images, scale bar = 100µm. B-H: Data are displayed in mean ± SD. I: Data are displayed in mean ± SEM. *P < 0.05, **P < 0.01, ***P < 0.001 and ****P < 0.0001 versus NM; #P < 0.05 versus CM; $$P < 0.01 versus Db/+.

In C57BL/6 mouse aortic rings, canagliflozin significantly reduced calcium deposition at 1.25 and 5 µM (**Fig. 3B**). Empagliflozin reduced calcification at 1.25 µM, but this effect was not observed at 5 µM, whereas dapagliflozin had no significant effect (**Fig. 3B**). In rat aortic rings, canagliflozin also reduced medial calcification, while dapagliflozin and empagliflozin did not significantly affect von Kossa staining (**Fig. 3C**).

To extend these findings to more complex biological and pathological conditions, we investigated vascular tissues from diabetic mice and humans. First, we examined aortic rings from diabetic db/db mice and non-diabetic db/+ controls. Under calcifying conditions, db/db aortic rings exhibited greater calcium deposition than db/+ rings, while canagliflozin reduced calcification in both genotypes (**Fig. 3E**). Importantly, to assess the translational relevance of these findings, we extended our investigations to human vascular explants from multiple arterial beds, including aortic patches, resistance arteries, iliac arteries, and epigastric arteries. Canagliflozin consistently reduced calcium deposition across these human vascular tissues, whereas dapagliflozin and empagliflozin showed no significant effect (**Fig. 3D**). Together, these findings establish that the direct anti-calcifying activity of canagliflozin extends beyond conventional experimental models and is conserved across multiple human vascular beds, supporting its translational relevance.

Similar results were obtained in primary hASMCs and MOVAS cells, in which canagliflozin, but not dapagliflozin or empagliflozin, reduced calcium deposition under calcifying conditions (**Fig. 3F,G**). In MOVAS cells, canagliflozin also dose-dependently reduced mRNA expression of the osteogenic markers *Spp1* and *Runx2*, whereas dapagliflozin and empagliflozin had no significant effect (**Fig. 3H** and **Supplementary Fig. S3**). To further assess the concentration-dependent effects of gliflozins in MOVAS cells, calcification was analyzed by OsteoSense staining after treatment with canagliflozin, dapagliflozin or empagliflozin at 1.25, 2.5, 5 or 10 µM. Canagliflozin reduced the OsteoSense signal at all tested concentrations, whereas dapagliflozin and empagliflozin did not modify calcification at any dose (**Fig. 3I**). At 2.5, 5 and 10 µM, canagliflozin also reduced total cell number (**Fig. 3I**), suggesting a possible effect of canaglifozin on cell expansion and/or viability.

### Canagliflozin reduces MOVAS cell proliferation and migration under calcifying conditions

To further investigate the cellular mechanisms associated with the anti-calcifying effect of canagliflozin, we assessed key vascular smooth muscle cell processes involved in calcifying remodeling, including proliferation, migration and cell viability. MOVAS cell proliferation and migration were monitored in real time using the IncuCyte system. Calcification medium (CM) did not significantly affect MOVAS cell proliferation compared with normal medium (NM) (**Fig. 4A**). Under CM conditions, canagliflozin reduced MOVAS cell proliferation at all tested concentrations, whereas dapagliflozin and empagliflozin had no significant effect (**Fig. 4B**). In scratch-wound assays, CM accelerated wound closure compared with NM, as reflected by increased relative wound density over time (**Fig. 4C**). Under CM conditions, canagliflozin reduced MOVAS cell migration at 10 µM, while dapagliflozin and empagliflozin did not significantly alter wound closure (**Fig. 4D**).

**Figure 4.**
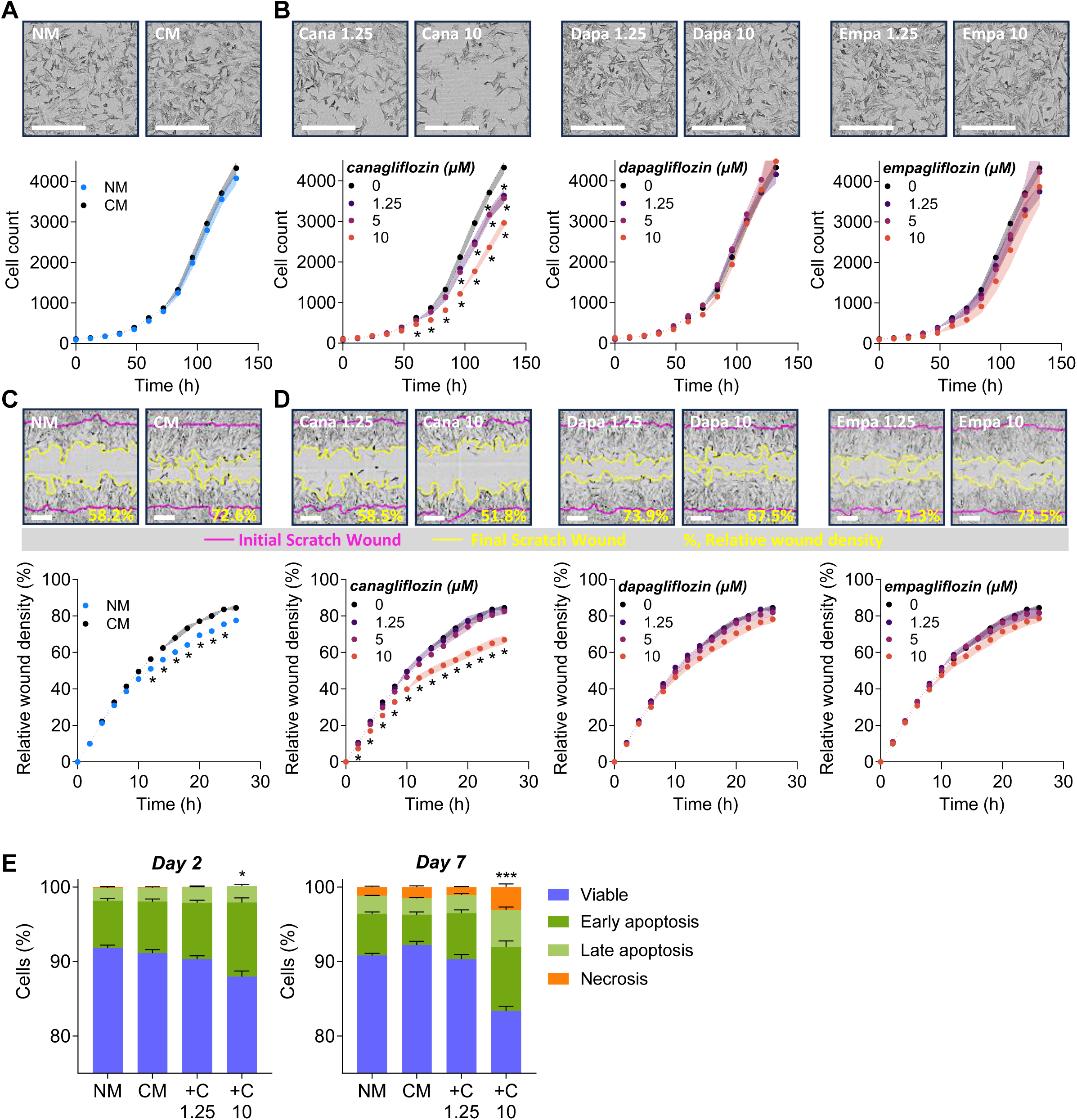
Canagliflozin alters MOVAS proliferation, migration and cell survival under calcifying conditions. **(A, B)** MOVAS cell proliferation was monitored in real time using the IncuCyte live-cell imaging system. **(A)** Proliferation curves and representative images of MOVAS cells cultured in normal medium (NM) or calcification medium (CM), and **(B)** in CM with canagliflozin, dapagliflozin or empagliflozin at 0, 1.25, 5 or 10 µM. Scale bar = 400µm. **(C, D)** MOVAS cell migration was assessed using an IncuCyte scratch-wound assay and quantified as relative wound density (%). **(C)** Time-course analysis and representative images of wound closure in MOVAS cells cultured in NM or CM, and **(D)** in CM with canagliflozin, dapagliflozin or empagliflozin at 0, 1.25, 5 or 10 µM. Scale bar = 200µm. **(E)** Cell viability and death were assessed by IncuCyte Annexin V/PI analysis after 2 days and 7 days of treatment with NM, CM, or CM with canagliflozin at 1.25 or 10 µM (CM+C1.25 and CM+C10, respectively). Stacked plots show the distribution of viable cells (Annexin V−/PI−), early apoptotic cells (Annexin V+/PI−), late apoptotic cells (Annexin V+/PI+) and necrotic cells (Annexin V−/PI+). Statistical analysis was performed on the combined apoptotic/damaged fraction, defined as early apoptotic + late apoptotic + necrotic cells. Data are displayed in mean ± SEM. *P < 0.05 versus CM.

Cell viability and death were then assessed by Annexin V/PI staining after 2 and 7 days of treatment (**Fig. 4E**). At 1.25 µM, a concentration at which canagliflozin reduced calcification and proliferation, canagliflozin did not modify the combined apoptotic/damaged fraction, defined as early apoptotic, late apoptotic and necrotic cells, compared with CM. In contrast, 10 µM canagliflozin increased the proportion of apoptotic/damaged cells, indicating that cell death may contribute to the reduced cell expansion observed at higher concentrations, but does not account for the anti-proliferative effect observed at lower doses.

### Canagliflozin reduces glucose uptake and mitochondrial ATP production in MOVAS cells

Given the effects of canagliflozin on MOVAS proliferation and migration, we next investigated whether these effects were accompanied by alterations in glucose uptake and cellular bioenergetics. Radiolabeled 2-deoxyglucose uptake assays showed that canagliflozin dose-dependently reduced glucose uptake, whereas dapagliflozin and empagliflozin had no significant effect (**Fig. 5A**).

**Figure 5.**
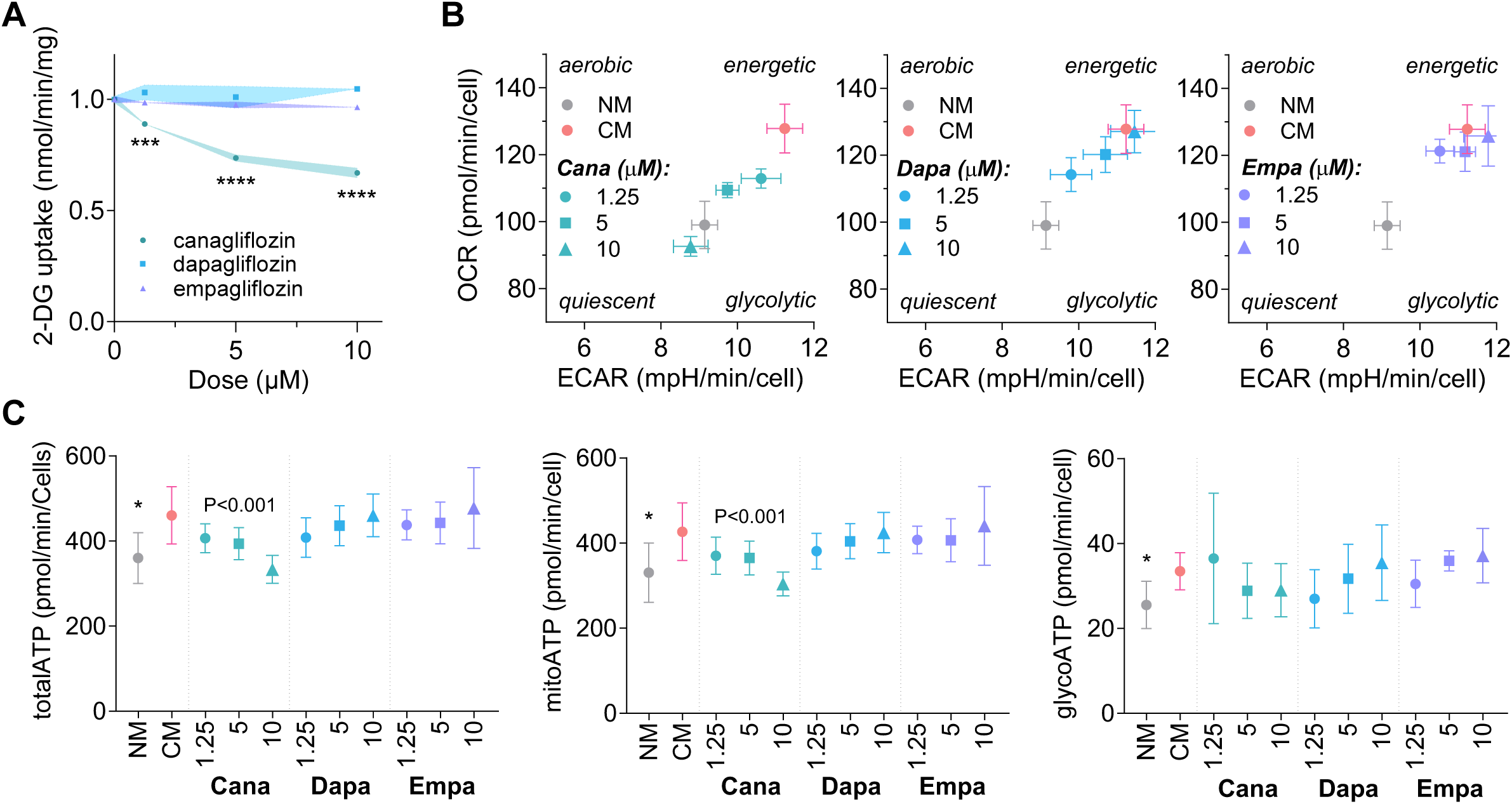
Effects of gliflozins on glucose uptake and cellular bioenergetics in MOVAS cells. **(A)** Radiolabeled 2-deoxyglucose uptake in MOVAS cells cultured with canagliflozin, dapagliflozin or empagliflozin at 0, 1.25, 5 or 10 µM. **(B–C)** Seahorse ATP rate analysis in MOVAS cells cultured in normal medium (NM), calcification medium (CM), or CM treated with canagliflozin, dapagliflozin or empagliflozin (1.25, 5 or 10 µM) for 2 days. **(B)** Bioenergetic maps showing basal oxygen consumption rate (OCR) and extracellular acidification rate (ECAR). **(C)** Total, mitochondrial and glycolytic ATP production rates. A: Data are displayed in mean ± SEM. B: Data are displayed in mean ± SD. *P < 0.05 versus CM.

We next examined whether these differential effects were accompanied by changes in cellular bioenergetics using the Seahorse ATP rate assay after 2 days of treatment. CM increased both extracellular acidification rate (ECAR) and oxygen consumption rate (OCR) compared with NM, indicating increased bioenergetic activity under calcifying conditions (**Fig. 5B**). Canagliflozin reduced ECAR and OCR in a dose-dependent manner compared with CM, whereas dapagliflozin and empagliflozin had no significant effect at any tested concentration (**Fig. 5B**). ATP rate analysis further showed that CM increased total ATP production, with increases in both glycolytic and mitochondrial ATP production (**Fig. 5C**). Canagliflozin dose-dependently reduced total ATP production, primarily reflecting a decrease in mitochondrial ATP production without significantly affecting glycolytic ATP production (**Fig. 5C**). Dapagliflozin and empagliflozin did not significantly alter any of these parameters.

### SGLT2 and SGLT1 expression is minimal in vascular tissues and smooth muscle cells

Given the selective effects of canagliflozin on vascular calcification, proliferative and migratory responses, bioenergetic activity and glucose uptake, we next examined whether SGLT2 or SGLT1 expression could account for this profile in vascular tissue and cells. Western blot analysis using a SGLT2 knockout-validated antibody^42^ detected SGLT2 in wild-type mouse kidney but not in SGLT2-knockout kidney, confirming antibody specificity, whereas negligible SGLT2 signal was detected in aorta or MOVAS cells (**Fig. 6A**). Consistently, immunofluorescence showed strong SGLT2 staining in the kidney, but minimal staining in aorta and MOVAS cells (**Fig. 6B**). SGLT1 showed a similar expression pattern, detected in mouse kidney but minimal staining in aorta and MOVAS cells.

**Figure 6.**
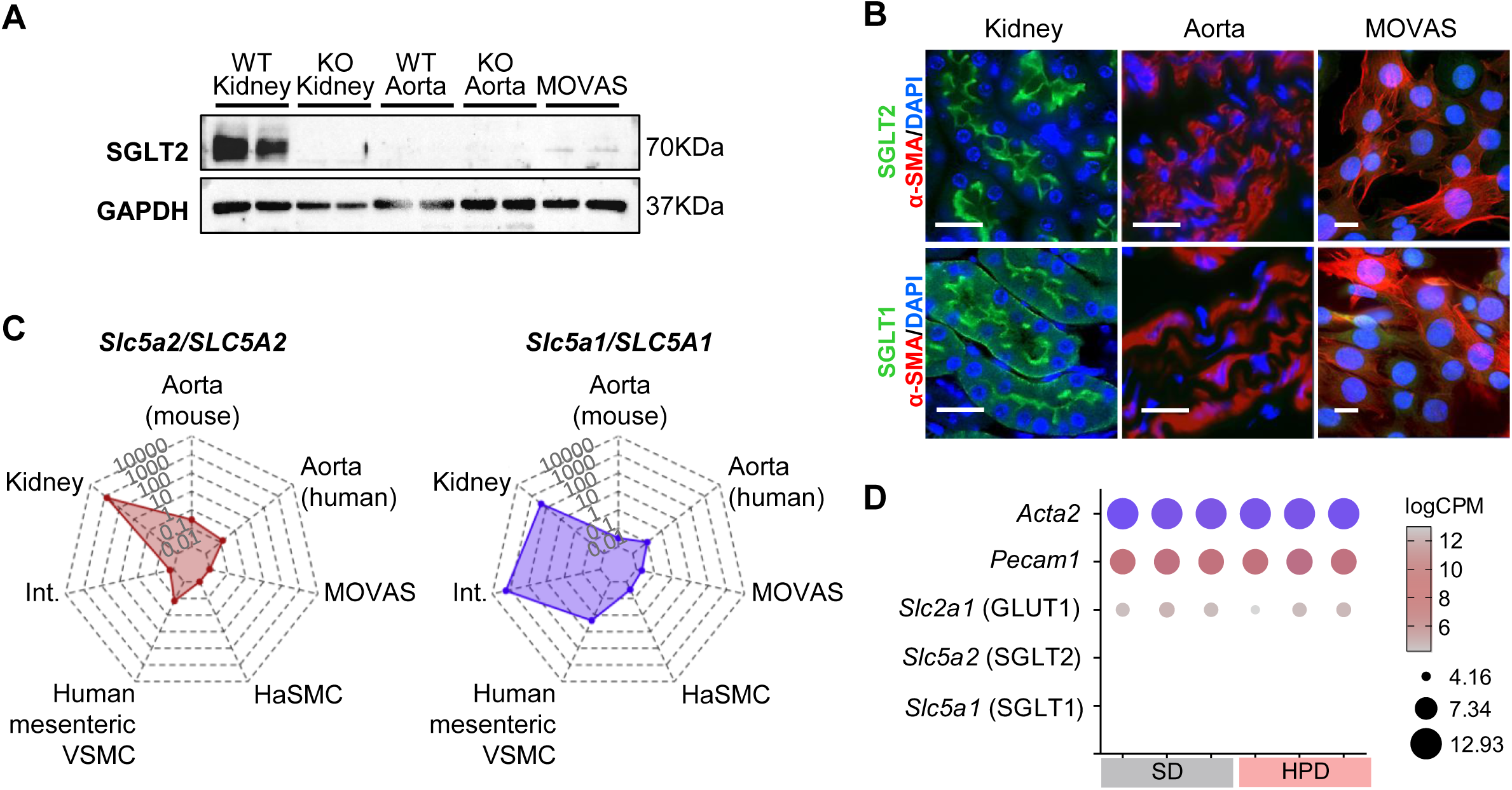
SGLT2 and SGLT1 expression in vascular tissues and smooth muscle cells. **(A)** Western blot analysis of SGLT2 protein expression in wild-type (WT) and SGLT2-knockout (KO) mouse kidneys, mouse aorta and MOVAS cells, using an SGLT2 knockout-validated antibody. **(B)** Representative immunofluorescence images showing SGLT2 and SGLT1 protein expression in mouse kidney, aorta and MOVAS cells. Scale bar = 20 µm. **(C)** TaqMan RT-qPCR analysis of *Slc5a2/SLC5A2* and *Slc5a1/SLC5A1* expression in mouse kidney, intestine (Int.) and aorta, in human aorta, and in vascular smooth muscle cells, including MOVAS cells, primary human aortic smooth muscle cells (hASMCs) and human mesenteric artery-derived VSMCs. **(D)** Aortic RNA-seq analysis of mice fed a standard diet (SD) or high-phosphate diet (HPD).

TaqMan RT-qPCR confirmed robust kidney expression of *Slc5a2/SLC5A2* encoding SGLT2, and expression of *Slc5a1/SLC5A1* encoding SGLT1 in both kidney and intestine, while both transporters were weakly expressed or undetectable in mouse aorta, human aorta, MOVAS cells, primary hASMCs and human mesenteric artery-derived VSMCs (**Fig. 6C**). Analysis of the Tabula Muris and Tabula Sapiens single-cell RNA-seq atlases further supported the low or undetectable expression of both transporters in aortic cell populations (**Supplementary Fig. S4**).

Given that HPD induced inflammatory and cellular stress responses in the aorta (**Fig.2D**), we next investigated whether this pro-calcifying environment could also induce vascular SGLT2 or SGLT1 expression. Analysis of aortic RNA-seq data from SD- and HPD-fed mice revealed detectable expression of vascular markers and the glucose transporter GLUT1 (*Slc2a1*), whereas neither *Slc5a1* nor *Slc5a2* was retained in the dataset (**Fig. 5D**).

Together, these findings indicate that canagliflozin exerts its vascular effects in a setting of minimal SGLT1 and SGLT2 expression, consistent with the involvement of mechanisms beyond canonical vascular SGLT inhibition.

### Sotagliflozin and mizagliflozin do not reproduce the anti-calcifying effect of canagliflozin

To further test whether sodium-glucose cotransporter inhibition could explain the anti-calcifying effect of canagliflozin, we evaluated sotagliflozin, a dual SGLT1/2 inhibitor, and mizagliflozin, an SGLT1 inhibitor, in the DBA/2N HPD model. HPD mice were treated with sotagliflozin or mizagliflozin at 3 mg/kg/day. At this dose, sotagliflozin induced urinary glucose excretion, confirming systemic sodium-glucose cotransporter inhibition (**Supplementary Fig. S5)**. In contrast, mizagliflozin did not induce glycosuria, indicating limited systemic effect at this dose and limiting the interpretation of its *in vivo* effect (**Supplementary Fig. S5).** Neither sotagliflozin nor mizagliflozin significantly reduced aortic calcification in HPD mice (**Fig. 7A**).

**Figure 7.**
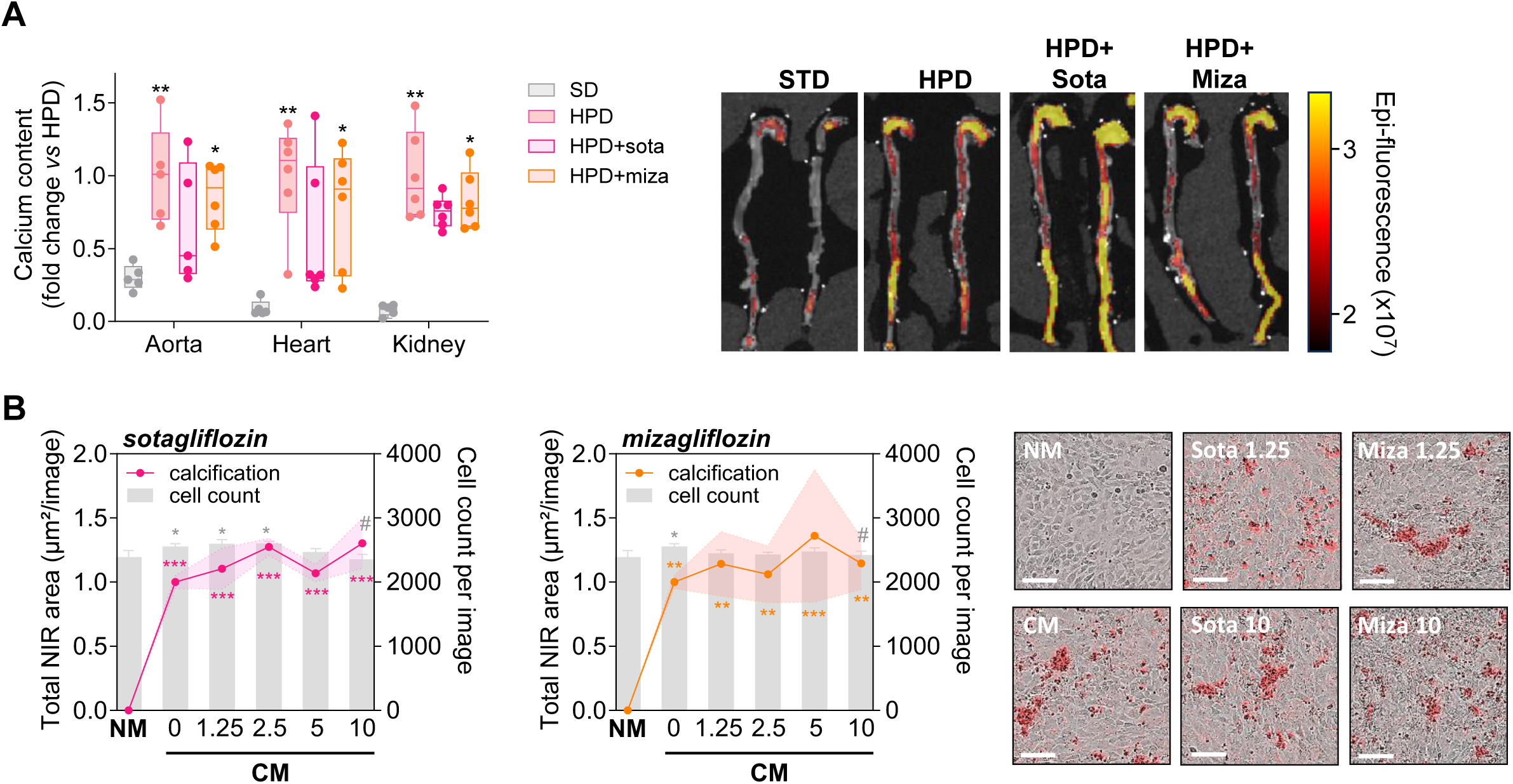
Sotagliflozin and mizagliflozin do not reproduce the anti-calcifying effect of canagliflozin. **(A)** Calcium content in the aortic arch, heart and kidney of DBA/2N mice fed a standard diet (SD) or high-phosphate diet (HPD) for 2 weeks and treated with sotagliflozin or mizagliflozin (3 mg/kg/day), with representative IVIS fluorescence images of aortas following OsteoSense administration. Data are displayed in mean ± SD. *P < 0.05 and **P < 0.01 versus SD. **(B)** Dose-response analysis of MOVAS calcification under calcifying conditions after treatment with sotagliflozin or mizagliflozin (1.25, 2.5, 5 or 10 µM), assessed by OsteoSense staining, with quantification and representative images, scale bar = 100µm. Data are displayed in mean ± SEM. *P < 0.05, **P < 0.01, ***P < 0.001 and ****P < 0.0001 versus NM.

To directly assess these compounds in vascular cells independently of systemic exposure, sotagliflozin and mizagliflozin were then tested in MOVAS cells using OsteoSense dose-response assay. Neither compound reduced MOVAS calcification at any tested concentration (**Fig. 7B**). The lack of effect of sotagliflozin and mizagliflozin in MOVAS cells, together with the low expression of SGLT2 and SGLT1 in vascular tissues and cells, indicates that pharmacological SGLT1/2 or SGLT1 inhibition is not sufficient to explain the anti-calcifying effect of canagliflozin.

## Discussion

Vascular calcification is a major contributor to cardiovascular morbidity and mortality in patients with CKD, yet effective treatments remain limited.^43^ Although SGLT2i have substantially improved cardiorenal outcomes, their effects on vascular calcification and the mechanisms underlying their direct vascular actions remain incompletely understood. Here, we show that canagliflozin, unlike dapagliflozin and empagliflozin, prevents phosphate-induced vascular calcification across complementary experimental models. Beyond its anti-calcifying activity, our findings suggest that canagliflozin may interfere with the phenotypic and metabolic activation of VSMC, through mechanisms that are not readily explained by inhibition of their canonical SGLT targets.

The differential effects of the three gliflozins are particularly interesting given their shared primary pharmacological action. Importantly, insufficient dosing is unlikely to fully explain the absence of anti-calcifying effects with dapagliflozin and empagliflozin. First, all three compounds induced glycosuria in vivo, confirming their pharmacological activity. The administered dose (3 mg/kg/day) corresponds to a human equivalent dose of approximately 15-28 mg/day, within the range of clinically prescribed doses for dapagliflozin and empagliflozin and below those used for canagliflozin.^29,30,44^ Second, *ex vivo* and *in vitro*, dapagliflozin and empagliflozin failed to reproduce the anti-calcifying effect of canagliflozin even at concentrations up to 10 µM. The concentrations tested *in vitro* (1.25–10 µM) were within or slightly above the range of peak total plasma concentrations achieved with therapeutic doses of canagliflozin (approximately 2.7 µM at 100 mg/day and 10.5 µM at 300 mg/day), whereas they exceeded those reported for dapagliflozin (approximately 0.4 µM at 10 mg/day) and empagliflozin (approximately 0.7 µM at 25 mg/day).^45–48^ Nevertheless, given the high plasma protein binding of these compounds, particularly canagliflozin, direct extrapolation of *in vitro* concentrations to clinically relevant free drug exposure remains challenging.^29,30^ Although the culture medium contained FBS, its lower protein content compared with human plasma may result in higher unbound drug fractions. Furthermore, differences in pharmacokinetics, tissue exposure, and off-target potency must be considered when interpreting these compound-specific effects, particularly if they involve mechanisms beyond canonical SGLT2 inhibition.

The bone-vascular calcification paradox, characterized by the coexistence of ectopic vascular mineral deposition and impaired skeletal mineralization, is a well-recognized feature of CKD mineral and bone disorder. This relationship raises the question of whether therapies targeting vascular calcification might also affect physiological bone remodeling. ^49,50^ In this context, we investigated whether the anti-calcifying effects of canagliflozin were accompanied by alterations in bone remodeling. Although HPD increased indices of both bone formation and resorption, none of the gliflozins significantly affected bone histomorphometric parameters. Clinical evidence regarding the skeletal effects of SGLT2i remains mixed, particularly for canagliflozin, with inconsistent findings on bone mineral density and fracture risk across clinical trials.^51–53^ In contrast, dapagliflozin and empagliflozin have generally shown no significant adverse skeletal effects. Preclinical studies have also reported conflicting skeletal effects of canagliflozin.^54–57^ Thus, while our findings suggest that canagliflozin attenuates pathological vascular mineralization without detectable short-term alterations in bone remodeling, the limited duration of our model precludes conclusions regarding long-term skeletal effects. Whether canagliflozin similarly affects mitochondrial ATP production in osteoblasts, as observed in VSMCs in our study, and potentially influences bone formation remains to be investigated.

One possible interpretation of our findings is that canagliflozin limits the activation of VSMCs. Under physiological conditions, differentiated VSMCs maintain a predominantly quiescent, contractile phenotype. In response to phosphate overload, inflammation, or vascular injury, they undergo phenotypic remodeling and may acquire proliferative, migratory, inflammatory, or osteogenic characteristics. These responses are not necessarily sequential but represent interconnected programs contributing to vascular remodeling. Thus, proliferation and migration are not synonymous with osteogenic differentiation, yet they reflect aspects of VSMC activation that may facilitate the establishment of a pro-calcifying environment.^58,59^ In this context, our findings are consistent with those of Behnammanesh et al. (2020), who demonstrated that canagliflozin inhibits VSMC proliferation and migration, induces G₀/G₁ cell-cycle arrest, and promotes HO-1 expression, with effects not reproduced by dapagliflozin or empagliflozin under comparable experimental conditions.^31^ Our study extends these observations to vascular calcification, suggesting that limiting VSMC activation may contribute to the anti-calcifying activity of canagliflozin.

The metabolic effects of canagliflozin may provide a mechanistic link between these cellular responses. VSMC phenotypic switching requires substantial metabolic adaptations to sustain proliferation, migration, and extracellular matrix remodeling. In our experiments, canagliflozin reduced glucose uptake and total ATP production, primarily through decreased mitochondrial ATP production, potentially limiting the energetic and biosynthetic demands of an activated phenotype. Interestingly, canagliflozin has previously been shown to inhibit mitochondrial complex I and activate AMPK independently of SGLT2 inhibition.^60^ Rather than acting directly on osteogenic differentiation or mineral deposition, canagliflozin may affect the energy-dependent processes underlying VSMC activation and downstream pathological vascular remodeling. However, whether these metabolic alterations contribute causally to its anti-calcifying activity remains to be established.

Our aortic transcriptomic findings provide complementary support for the involvement of these processes. Although several endothelial and inflammatory responses were shared among gliflozins, canagliflozin additionally reversed HPD-induced alterations in genes associated with osteogenic signaling, extracellular matrix remodeling, and cell growth. Notably, ATF4 promotes vascular calcification by increasing phosphate uptake through the regulation of PiT1 and PiT2, while TWEAK signaling through its receptor Fn14, encoded by *Tnfrsf12a*, enhances phosphate-induced VSMC osteogenic differentiation and calcification via NF-κB activation.^61,62^ Furthermore, cathepsin K, encoded by *Ctsk*, contributes to elastin degradation, generating matrix alterations and elastin-derived peptides that promote vascular mineralization.^63^ Changes in genes involved in nucleotide synthesis, such as *Ctps* and *Impdh2*, and mitochondrial metabolic regulators are also consistent with modulation of cellular growth and metabolic associated processes. The reversal of these transcriptional changes by canagliflozin may therefore reflect modulation of multiple convergent pro-calcifying mechanisms linking VSMC activation, metabolic regulation, and calcification.

The anti-calcifying effect of canagliflozin, which was not reproduced by other SGLT2 inhibitors (dapagliflozin and empagliflozin), the selective SGLT1 inhibitor mizagliflozin, or the dual SGLT1/SGLT2 inhibitor sotagliflozin, suggests that its direct vascular actions may involve mechanisms beyond canonical SGLT inhibition. The expression of SGLT2 outside the kidney remains controversial. While some studies have reported vascular SGLT2 expression, including under inflammatory conditions, others have found minimal or undetectable expression in cardiovascular tissues.^64–66^ These discrepancies may partly reflect methodological challenges, particularly the influence of tissue fixation on protein detection and the limited specificity of certain anti-SGLT2 antibodies.^67,68^ To address these issues, we combined complementary approaches, including transcriptomic and RT-qPCR analyses, immunofluorescence, and Western blotting using an SGLT2-knockout-validated antibody. All consistently indicated minimal or undetectable SGLT2 expression in vascular tissues and VSMCs. Importantly, despite the pronounced inflammatory and cellular stress responses induced by HPD, neither *Slc5a2* nor *Slc5a1* was present in the aortic RNA-seq dataset, providing no evidence of substantial induction under pro-calcifying conditions.

Our findings are consistent with previous reports suggesting SGLT2-independent actions of gliflozins. For canagliflozin, these include modulation of mitochondrial function and AMPK signaling as well as direct anti-inflammatory effects in vascular endothelial cells lacking detectable *SLC5A2* transcripts.^69,70^ Similarly, studies using SGLT2-knockout mice have demonstrated that empagliflozin retains its cardioprotective effects against myocardial ischemia-reperfusion injury and heart failure, supporting SGLT2-independent mechanisms in distinct cardiovascular settings.^71,72^ Together, these findings support the involvement of noncanonical mechanisms in the vascular effects of gliflozins, although the precise molecular target mediating the anti-calcifying activity of canagliflozin remains to be identified.

A major strength of our study lies in the systematic comparison of three SGLT2 inhibitors across complementary *in vivo*, *ex vivo*, and *in vitro* models, together with dose-response experiments that strengthen the interpretation of their differential effects. To our knowledge, this study is the first to demonstrate a differential effect of gliflozins on VC, as the mechanisms and clinical benefits of SGLT2i have generally been considered to largely reflect a class effect rather than drug-specific properties. Importantly, the anti-calcifying activity of canagliflozin was confirmed in murine and human vascular explants, which preserve native vessel architecture, extracellular matrix, and interactions between resident cells, providing a physiologically relevant model beyond isolated VSMCs. Furthermore, its effectiveness in aortic rings from diabetic db/db mice suggests that this protection may also extend to a diabetic environment, where hyperglycemia, advanced glycation end products, and oxidative stress contribute to exacerbate vascular injury. Together, the consistency of these findings across experimental models and concentrations, including human tissues, strengthens their robustness and translational relevance.

Nevertheless, several limitations remain. Although our findings suggest that canagliflozin attenuates VSMC activation through metabolic and phenotypic changes, the molecular target and causal mechanisms underlying its anti-calcifying activity remain to be established. In addition, concentration-dependent effects on cell viability warrant caution when extrapolating these findings to clinical settings. Furthermore, in our *in vivo* study, none of the gliflozins significantly improved renal function, despite their established nephroprotective effects in CKD patients. However, these benefits on GFR are primarily a long-term effect which may not be captured by our short and severe nephrocalcinosis model.^16,18^

Another limitation concerns discrepancies with previous studies reporting anti-calcifying effects of dapagliflozin and empagliflozin.^23–25,73–75^ These studies differ substantially in genetic background, sex, treatment duration, and calcification-inducing stimuli, including diabetic and atherosclerotic conditions, vitamin D3 overload, or subtotal nephrectomy combined with phosphate loading. Such differences may influence both the predominant calcification mechanisms and treatment responses. Notably, some studies used male C57BL/6 mice, a strain relatively resistant to vascular calcification, whereas our model employed female DBA/2N mice, which have a genetic background more susceptible to ectopic mineralization.^76^ Furthermore, whereas our model is predominantly driven by severe phosphate-induced mineral disturbances, diabetic and atherosclerotic models involve additional mechanisms, including hyperglycemia, oxidative stress, and inflammation. Interestingly, even within similar models, conflicting results have been reported. Li et al. (2023) demonstrated that empagliflozin reduced atherosclerotic calcification in male ApoE-deficient mice, whereas Kalanski et al. (2024) found no reduction in total aortic calcium accumulation in females, further suggesting that anti-calcifying responses to SGLT2i may depend on the experimental context.^27,74^ Thus, rather than challenging the established cardiorenal class effects of SGLT2i, our findings suggest that their anti-calcifying effects may depend on both the individual compound and the underlying pathological context. This concept could ultimately support a more individualized use of SGLT2i according to the underlying disease phenotype and the mechanisms predominantly involved.

Our findings challenge the assumption that all gliflozins exert identical direct vascular effects and identify canagliflozin as a candidate modulator of phosphate-induced vascular calcification. Beyond reducing mineral deposition canagliflozin is associated with metabolic and phenotypic changes consistent with reduced VSMC activation. Whether these compound-specific actions translate into clinically meaningful protection against vascular ageing and calcification in CKD remains to be established.

## Acknowledgments

We thank the Phenotyping Department of the UMS006 CREFRE-Anexplo platform and the following I2MC facilities (Inserm, Toulouse, France) for their technical support: GeT-Santé (GenoToul GeT), Phi and Cytometry (GenoToul TRI), We-Met (functional biochemistry), and Bioinformatics. We are also grateful to Awahan Rahman for technical assistance. We acknowledge the PerMediK COST Action (CA21165; Personalized Medicine in Chronic Kidney Disease: Improved Outcome Based on Big Data) and its consortium members. JMDT holds a contract funded by the European Union’s Horizon Europe research and innovation programme under the Marie Skłodowska-Curie Actions (MSCA), grant agreement No. 101208747, and received funding from the Spanish Society of Nephrology and the Carlos III Health Institute (ISCIII) through the projects PMP21/00109, EU-Calprotect (AC24/00005), and co-funded by the European Union. PS and KK were supported by the Swedish Heart and Lung Foundation, CIMED, ALF and Westman foundation. PS and JPS are members of EUTox.

## Data availability

The RNA-seq data generated in this study will be deposited in the Gene Expression Omnibus (GEO) database and made publicly available upon publication of the manuscript. In addition, publicly available datasets used in the analyses were obtained from the Tabula Sapiens (https://tabula-sapiens.sf.czbiohub.org/) and Tabula Muris (https://www.czbiohub.org/sf/tabula-muris/) atlases.

## Supplementary figure legends

**Supplementary figure S1. Validation of SGLT inhibition by gliflozins in diabetic db/db and DBA/2N mice. (A, B)** Percentage change in blood glucose and urinary glucose excretion after treatment with canagliflozin, dapagliflozin or empagliflozin at 3 mg/kg/day in **(A)** db/db diabetic mice and **(B)** female DBA/2N mice. Values are expressed relative to the corresponding vehicle-treated control group. Data are shown as mean ± SD. *P < 0.05 versus vehicle.

**Supplementary figure S2. Gliflozin treatment does not modify renal injury, nephrocalcinosis or bone remodeling in HPD DBA/2N mice. (A)** RT-qPCR analysis of kidney markers *Havcr1* (KIM-1), *Adgre1* (F4/80), *Col3a1* and *Slc5a2* (SGLT2) in mice fed a standard diet (SD), a high-phosphate diet (HPD), or HPD combined with canagliflozin, dapagliflozin or empagliflozin. **(B-D)** Kidney immunostaining and mineralization analyses across the different experimental groups. **(B)** Quantification of immunostaining for F4/80, type III collagen and SGLT2. **(C)** Quantification of renal mineralization assessed by Alizarin Red staining. **(D)** Representative images of F4/80, type III collagen and SGLT2 immunostaining, and Alizarin Red staining. **(E)** Representative bone histology images of undecalcified distal femur sections stained with modified Masson-Goldner trichrome (blue arrows indicate osteoblasts; orange arrows indicate osteoclasts). Scale bar = 50 µm. Data are shown as mean ± SD. *P < 0.05, **P < 0.01, ***P < 0.001 and ****P < 0.0001 versus SD.

**Supplementary figure S3. Effects of dapagliflozin and empagliflozin on osteogenic marker expression in MOVAS cells.** Relative mRNA expression of *Spp1* and *Runx2* in MOVAS cells cultured in calcifying conditions and treated with dapagliflozin and empagliflozin (1.25 µM), assessed by RT-qPCR.

**Supplementary figure S4. Single-cell RNA-seq analysis of SGLT1 and SGLT2 expression in renal and aortic cell populations.** Expression of *Slc5a2/SLC5A2* (SGLT2) and *Slc5a1/SLC5A1* (SGLT1) across murine **(A)** and human **(B)** cell populations using the Tabula Muris and Tabula Sapien single-cell RNA-seq atlases, respectively.

**Supplementary figure S5. Sotagliflozin, but not mizagliflozin, induces glycosuria in DBA/2N mice.** Percentage change in blood glucose and urinary glucose excretion after treatment with sotagliflozin or mizagliflozin at 3 mg/kg/day in female DBA/2N mice. Values are expressed relative to the corresponding vehicle-treated control group. Data are shown as mean ± SD. **P < 0.01 versus vehicle.

