## Supplementary material for "Systematic comparison of gliflozins identifies a selective anti-calcifying effect of canagliflozin beyond canonical SGLT2 inhibition"

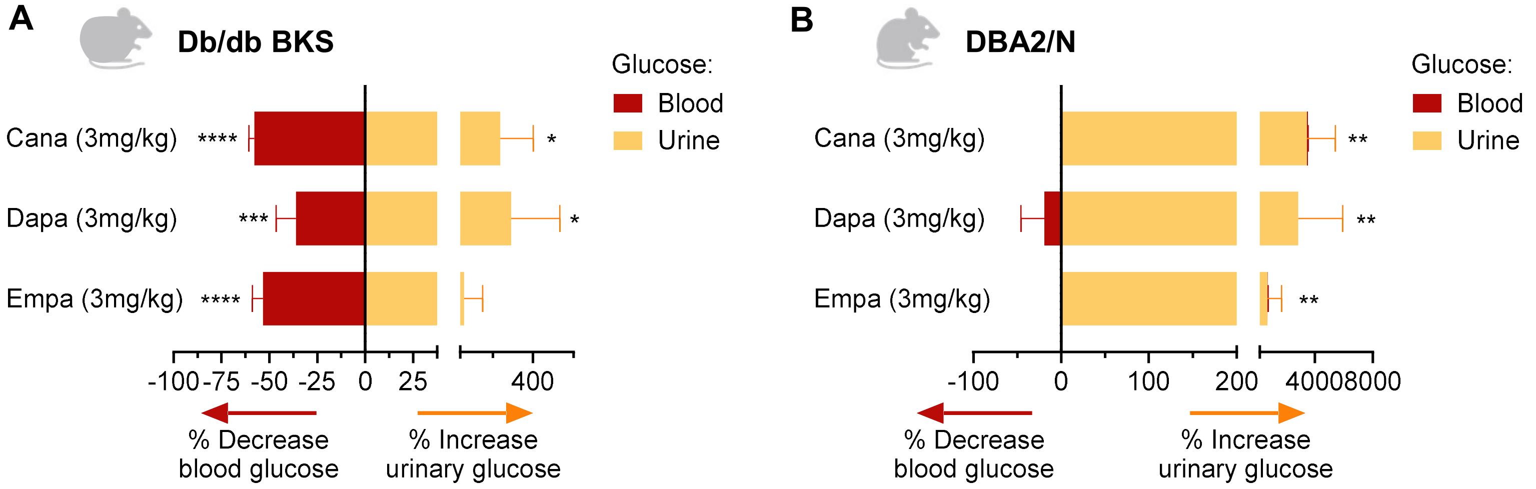


**Supplementary figure S1. Validation of SGLT inhibition by gliflozins in diabetic db/db and DBA/2N mice. (A, B)** Percentage change in blood glucose and urinary glucose excretion after treatment with canagliflozin, dapagliflozin or empagliflozin at 3 mg/kg/day in **(A)** db/db diabetic mice and **(B)** female DBA/2N mice. Values are expressed relative to the corresponding vehicle-treated control group. Data are shown as mean ± SD. *P < 0.05 versus vehicle.

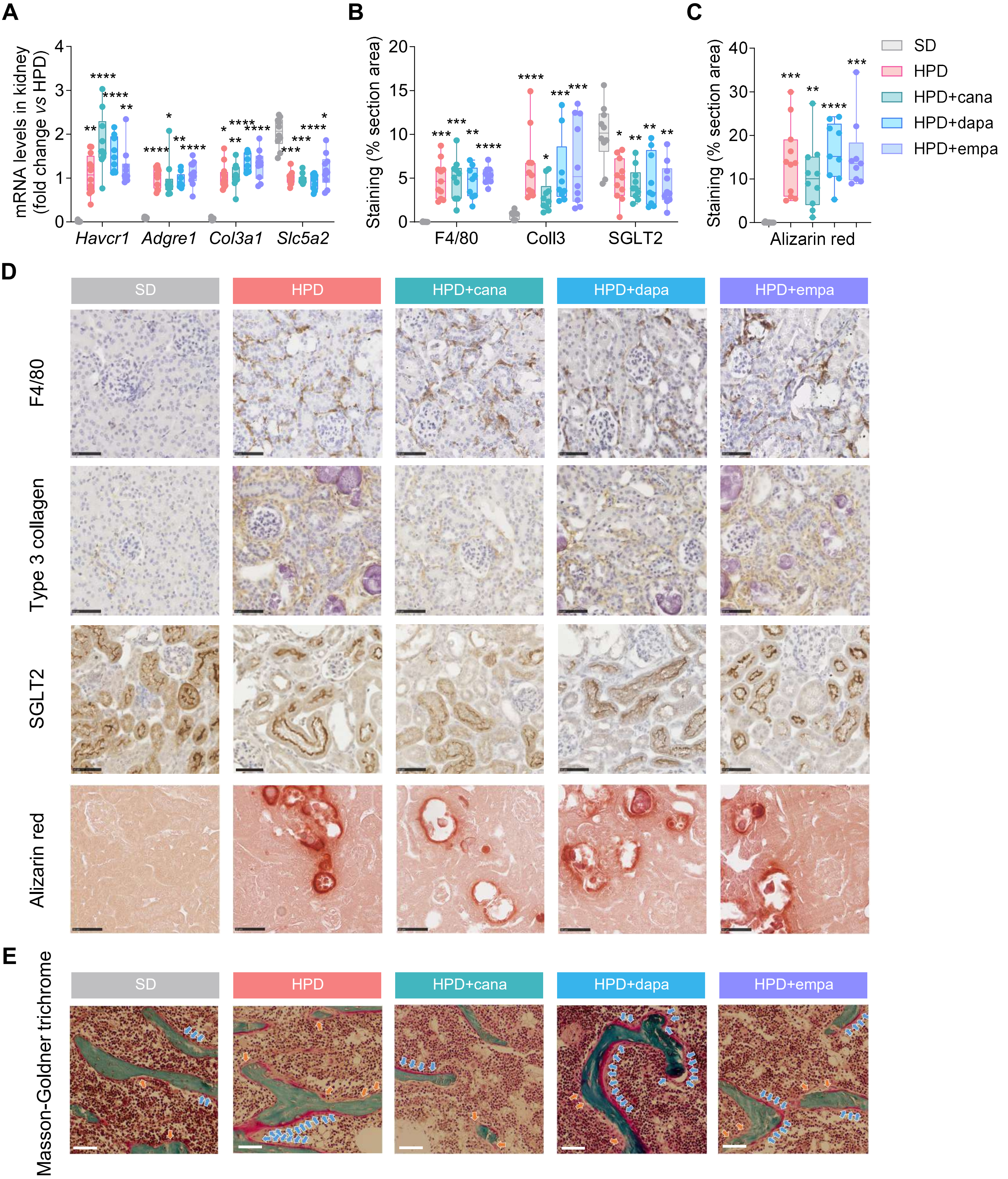


**Supplementary figure S2. Gliflozin treatment does not modify renal injury, nephrocalcinosis or bone remodeling in HPD DBA/2N mice. (A)** RT-qPCR analysis of kidney markers *Havcr1* (KIM-1), *Adgre1* (F4/80), *Col3a1* and *Slc5a2* (SGLT2) in mice fed a standard diet (SD), a high-phosphate diet (HPD), or HPD combined with canagliflozin, dapagliflozin or empagliflozin. **(B-D)** Kidney immunostaining and mineralization analyses across the different experimental groups. **(B)** Quantification of immunostaining for F4/80, type III collagen and SGLT2. **(C)** Quantification of renal mineralization assessed by Alizarin Red staining. **(D)** Representative images of F4/80, type III collagen and SGLT2 immunostaining, and Alizarin Red staining. **(E)** Representative bone histology images of undecalcified distal femur sections stained with modified Masson-Goldner trichrome (blue arrows indicate osteoblasts; orange arrows indicate osteoclasts). Scale bar = 50 µm. Data are shown as mean ± SD. *P < 0.05, **P < 0.01, ***P < 0.001 and ****P < 0.0001 versus SD.

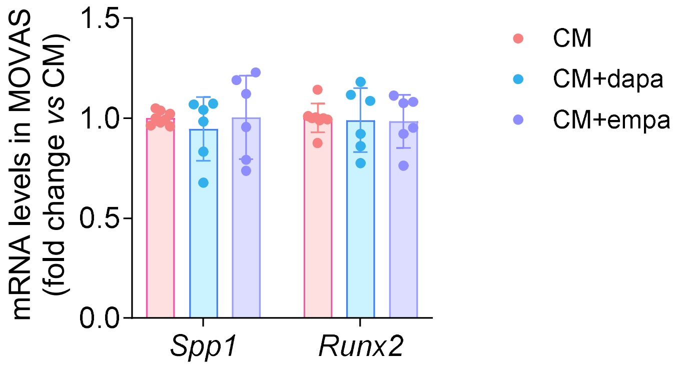


**Supplementary figure S3. Effects of dapagliflozin and empagliflozin on osteogenic marker expression in MOVAS cells.** Relative mRNA expression of *Spp1* and *Runx2* in MOVAS cells cultured in calcifying conditions and treated with dapagliflozin and empagliflozin (1.25 µM), assessed by RT-qPCR.

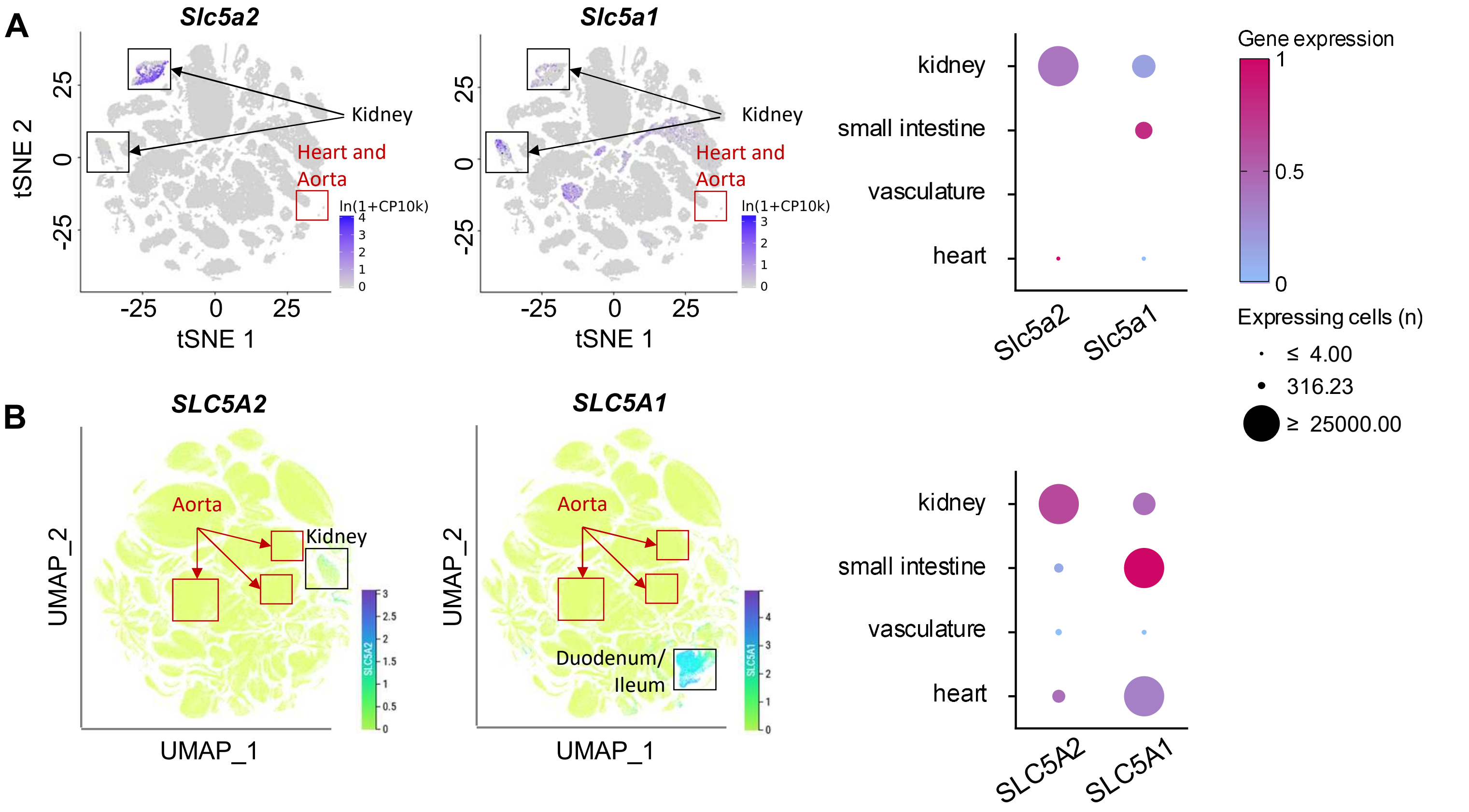


**Supplementary figure S4. Single-cell RNA-seq analysis of SGLT1 and SGLT2 expression in renal and aortic cell populations.** Expression of *Slc5a2/SLC5A2* (SGLT2) and *Slc5a1/SLC5A1* (SGLT1) across murine **(A)** and human **(B)** cell populations using the Tabula Muris and Tabula Sapien single-cell RNA-seq atlases, respectively.

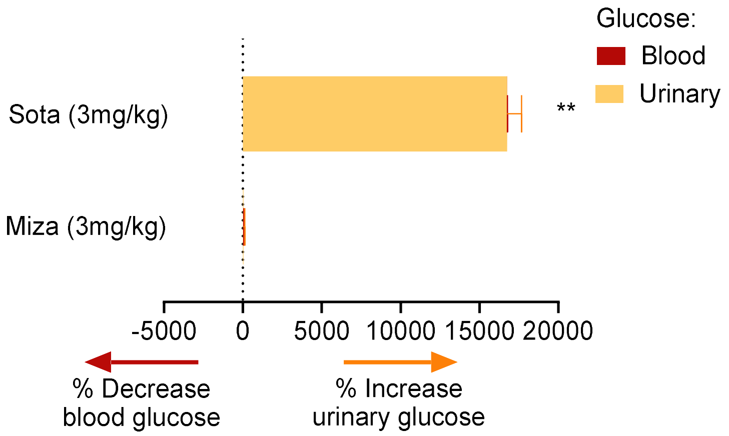


**Supplementary figure S5. Sotagliflozin, but not mizagliflozin, induces glycosuria in DBA/2N mice.** Percentage change in blood glucose and urinary glucose excretion after treatment with sotagliflozin or mizagliflozin at 3 mg/kg/day in female DBA/2N mice. Values are expressed relative to the corresponding vehicle-treated control group. Data are shown as mean ± SD. **P < 0.01 versus vehicle.
